# Is your data alignable? Principled and interpretable alignability testing and integration of single-cell data

**DOI:** 10.1101/2023.08.03.551836

**Authors:** Rong Ma, Eric D. Sun, David Donoho, James Zou

## Abstract

Single-cell data integration can provide a comprehensive molecular view of cells, and many algorithms have been developed to remove unwanted technical or biological variations and integrate heterogeneous single-cell datasets. Despite their wide usage, existing methods suffer from several fundamental limitations. In particular, we lack a rigorous statistical test for whether two high-dimensional single-cell datasets are alignable (and therefore should even be aligned). Moreover, popular methods can substantially distort the data during alignment, making the aligned data and downstream analysis difficult to interpret. To overcome these limitations, we present a spectral manifold alignment and inference (SMAI) framework, which enables principled and interpretable alignability testing and structure-preserving integration of single-cell data. SMAI provides a statistical test to robustly determine the alignability between datasets to avoid misleading inference, and is justified by high-dimensional statistical theory. On a diverse range of real and simulated benchmark datasets, it outperforms commonly used alignment methods. Moreover, we show that SMAI improves various downstream analyses such as identification of differentially expressed genes and imputation of single-cell spatial transcriptomics, providing further biological insights. SMAI’s interpretability also enables quantification and a deeper understanding of the sources of technical confounders in single-cell data.

## 1 Introduction

The rapid development of single-cell technologies has enabled the characterization of complex biological systems at unprecedented scale and resolution. On the one hand, diverse and heterogeneous single-cell datasets have been generated, enabling opportunities for integrative profiling of cell types and deeper understandings of the associated biological processes [83, 85, 79, 28]. On the other hand, the widely observed technical and biological variations across datasets also impose unique challenges to many downstream analyses [48, 81, 58]. These variations between datasets can originate from different experimental protocols, laboratory conditions, sequencing technologies, etc. It may also arise from biological variations when the samples come from distinct spatial locations, time points, tissues, organs, individuals, or species.

Several computational algorithms have been recently developed to remove the unwanted variations and integrate heterogeneous single-cell datasets. To date, the most widely used data integration methods, such as Seurat [77], LIGER [87], Harmony [44], fastMNN [35], and Scanorama [37], are built upon the key assumption that there is a shared latent low-dimensional structure between the datasets of interest. These methods attempt to obtain an alignment of the datasets by identifying and matching their respective low-dimensional structures. As a result, the methods would output some integrated cellular profiles, commonly represented as either a corrected feature matrix, or a joint low-dimensional embedding matrix, where the unwanted technical or biological variations between the datasets have been removed. These methods have played an indispensable role in current single-cell studies such as generating large-scale reference atlases of human organs [16, 61, 21, 94, 75], inferring lineage differentiation trajectories and pseudotime reconstruction [82, 68, 78], and multi-omic characterization of COVID-19 pathogenesis and immune response [76, 2, 47].

Despite their popularity, the existing integration methods also suffer from several fundamental limitations, which makes it difficult to statistically assess findings drawn from the aligned data. First, there is a lack of statistically rigorous methods to determine whether two or more datasets should be aligned. Without such a safeguard, existing methods are used to align and integrate single-cell datasets that do not have a meaningful shared structure, leading to problematic and misleading interpretations [17, 22]. Global assessment methods such as k-nearest neighbor batch effect test (kBET) [13], guided PCA (gPCA) [69], probabilistic principal component and covariates analysis (PPCCA) [65], and metrics such as local inverse Simpson’s index (LISI) [44], average silhouette width (ASW) [9], adjusted Rand index (ARI) [88], have been proposed to quantitatively characterize the quality of alignment or the extent of batch-effect removal, based on specific alignment procedures. However, these methods can only provide post-hoc evaluations of the mixing of batches, which may not necessarily reflect the actual alignability or structure-sharing between the original datasets. Moreover, these methods do not account for the noisiness (ASW, ARI, LISI) or the effects of high dimensionality (kBET, gPCA, PPCCA) of the single-cell datasets, resulting in biased estimates and test results. Other methods such as limma [70] and MAST [30] consider linear batch correction, whose focus is restricted to differential testing and does not account for batch effects causing shifts in covariance structures.

Moreover, research suggest that serious distortions to the individual datasets may be introduced by existing integration methods during their alignment process [17, 22]. In this study, we systematically evaluate the severity and effects of such distortions across several mostly used integration methods. Our results confirm that these methods, while eliminating the possible differences between datasets, may also alter the original biological signals contained in individual datasets, causing erroneous results such as power loss and false discoveries in downstream analyses. Finally, none of these popular integration methods admits a tractable closed-form expression of the final alignment function, with a clear geometric meaning of its constitutive components. As a result, these methods would suffer from a lack of interpretability, making it difficult to inspect and interpret the nature of removed variations, or to distinguish the unwanted variations from the potentially biologically informative variations.

To overcome the above limitations, we present a spectral manifold alignment and inference (SMAI) framework for accountable and interpretable integration of single-cell data. Our contribution is two-fold. First, we develop a rigorous statistical test (SMAI-test) that can robustly determine the alignability between two datasets. Secondly, motivated by this test, we propose an interpretable spectral manifold alignment algorithm (SMAI-align) that enables more reliable data integration without altering or corrupting the original biological signals. Our systematic experiments demonstrate that SMAI improves various downstream analyses such as the identification of cell types and their associated marker genes, and the prediction of single-cell spatial transcriptomics. Moreover, we show that SMAI’s interpretability provides insights into the nature of batch effects between single-cell datasets.

## 2 Results

### Overview of SMAI

SMAI consists of two components: SMAI-test flexibly determines the global or partial alignability between the datasets, whereas SMAI-align searches for the best similarity transformation to achieve the alignment. In particular, SMAI-test may be used in combination with existing integration methods, which only focus on the data alignment step.

SMAI-test evaluates the statistical significance against the null hypothesis that two single-cell datasets are (partially) alignable up to some similarity transformation, that is, combinations of scaling, translation, and rotation. It leverages random matrix theory to be robust to noisy, high-dimensional data. To increase flexibility, SMAI-test also allows for testing against partial alignability between the datasets, where the users can specify a threshold *t*%, so that the null hypothesis is that at least *t*% of the samples in both datasets are alignable. Recommended values for *t* are between 50 and 70 depending on the context, to ensure both sufficient sample size (or power), and flexibility to local heterogeneity (Methods); we used *t* = 60 for the real datasets analyzed in this study. Importantly, the statistical validity of SMAI-test is theoretically guaranteed over a wide range of settings (Methods, Theorem 4.1), especially suitable for modeling high-dimensional single-cell data. We support the empirical validity of the test with both simulated data and multiple real-world benchmark datasets, ranging from transcriptomics, chromatin accessibility, to spatial transcriptomics.

SMAI-align incorporates a high-dimensional shuffled Procrustes analysis, which iteratively searches for the sample correspondence and the best similarity transformation that minimizes the discrepancy between the intrinsic low-dimensional signal structures of the datasets. SMAI-align enjoys several advantages over the existing integration methods. First, SMAI-align returns an alignment function in terms of a similarity transformation, which has a closed-form expression with a clear geometric meaning of each component. The better interpretability enables quantitative characterization of the source and magnitude of any removed and remaining variations, and may bring insights into the mechanisms underlying the batch effects. Second, due to the shape-invariance property of similarity transformations, SMAI-align preserves the relative distances between the samples within individual datasets throughout the alignment, making the final integrated data less susceptible to technical distortions and therefore more suitable and reliable for downstream analyses. Third, unlike many existing methods (such as Seurat, Harmony and fastMNN), which require specifying a target dataset for alignment and whose performance is asymmetric with respect to the order of datasets, SMAI-align obtains a symmetric invertible alignment function that is indifferent to such an order, making its output more consistent and robust to technical artifacts.

### SMAI-test

Below we sketch the main ideas of the SMAI algorithm and leave the details to the Methods section. Suppose that 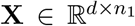 and 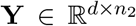 are the normalized count matrices generated from two single-cell experiments, with *d* being the number of features (genes) and *n*_1_ and *n*_2_ being the respective numbers of cells. To test for the alignability between **X** and **Y**, SMAI-test assumes a low-rank spiked covariance matrix model (Methods and Figure S1) where the low-dimensional signal structures of **X** and **Y** are encoded by the leading eigenvalues and eigenvectors of their corresponding population covariance matrices **Σ**_1_ and **Σ**_2_. As a result, the null hypothesis that the signal structures underlying **X** and **Y** are identical up to a similarity transformation implies that the leading eigenvalues of **Σ**_1_ and **Σ**_2_ are identical up to a global scaling factor. As such, a test statistic *T* (**X, Y**) based on comparing the leading eigenvalues of the empirical covariance matrices of **X** and **Y** can be computed, whose theoretical null distribution as (*n*_1_, *n*_2_, *d*) *→ ∞* can be derived precisely using random matrix theory. Thus, SMAI-test returns the p-value by comparing the test statistic *T* (**X, Y**) with its asymptotic null distribution (Figure 1a).

**Figure 1.**
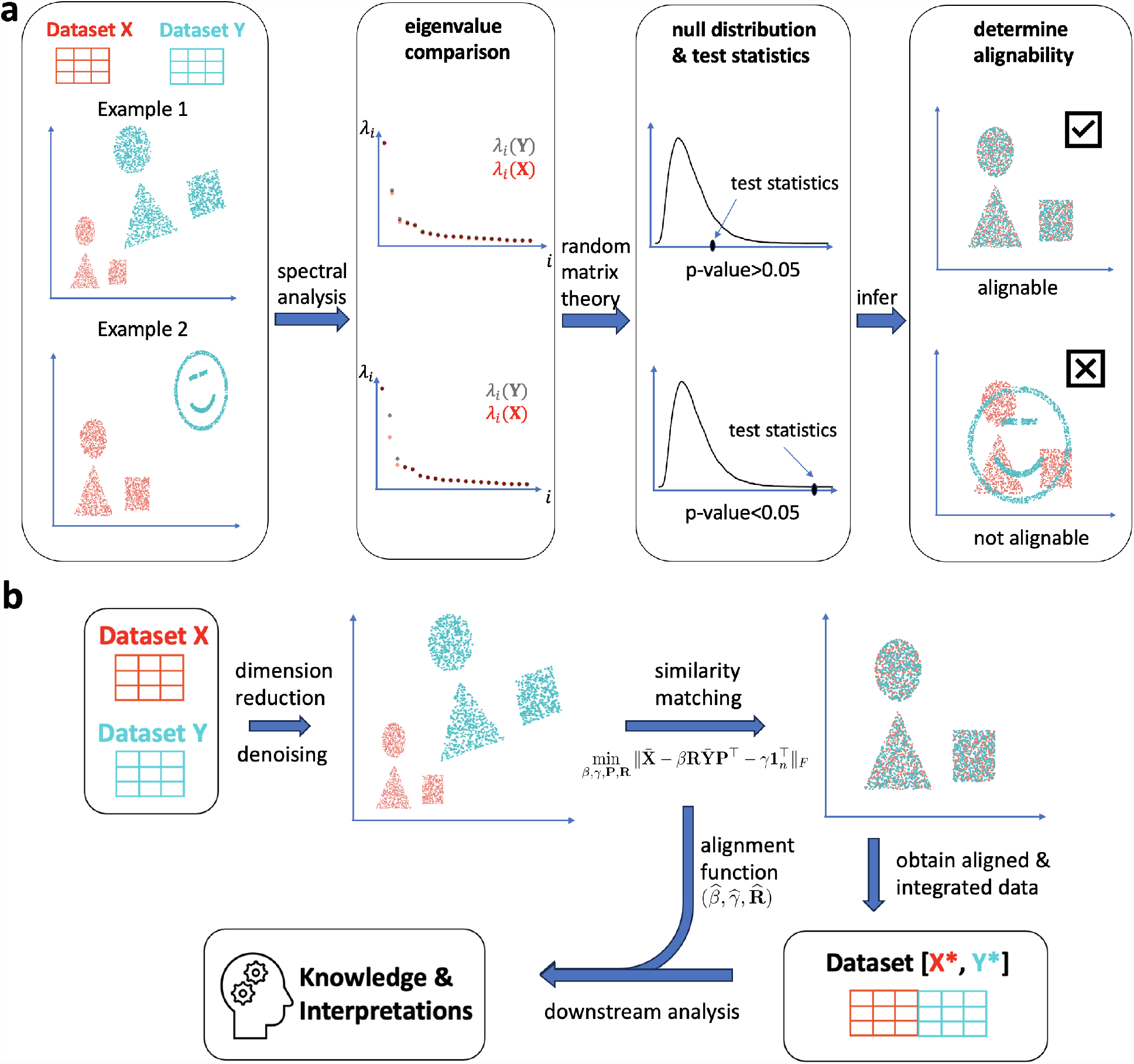
Overview and illustration of the SMAI algorithm. (a) SMAI-test imposes a low-rank spiked covariance matrix model where the low-dimensional signal structures of data matrices are encoded by a few largest eigenvalues of the population covariance matrices. Under the null hypothesis that the underlying signal structures are alignable up to a similarity transformation, a test statistic based on comparing the leading eigenvalues of the empirical covariance matrices is computed, whose theoretical null distribution as (*n, d*) *→ ∞* is derived using random matrix theory. The final p-value returned by SMAI-test is used to infer the alignability of the two datasets. (b) SMAI-align aims to solve for the shuffled Procrustes optimization problem (2.1). To do so, SMAI-align starts with a denoising procedure, and then adopts an iterative spectral algorithm to achieve similarity matching between the two datasets using high-dimensional Procrustes analysis. The method returns an integrated dataset containing all the samples with the original features, along with a closed-form alignment function, which is interpretable and can be readily used for various downstream analyses.

For the test of partial alignability, a sample splitting procedure is used where the first part is used to identify the subsets of each datasets with maximal correspondence or structure-sharing (Methods), and the second part is used to compute the test statistic and the p-value concerning the alignability between such maximal correspondence subsets. As such, the final p-value is free from selection bias or “double-dipping” due to repeated use of the samples in both selection and test steps.

### SMAI-align

SMAI-align starts by filtering out the low-rank signal structures in **X** and **Y** to obtain their denoised versions 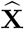 and Ŷ, and then approximately solves the following shuffled Procrustes optimization problem

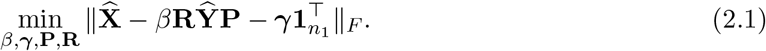

Here, **1**_*n*_ ∈ ℝ^*n*^ is an all-one vector, and the minimization is achieved for some global scaling factor *β* ∈ ℝ, some vector γ ∈ ℝ^*d*^ adjusting for the possible global mean shift between Ŷ and 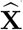,some extended orthogonal matrix (see Methods) 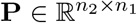 recovering the sample correspondence between 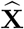 and Ŷ, and some rotation matrix **R** ∈ ℝ^*d×d*^ adjusting for the possible covariance shift. Compared with the traditional Procrustes analysis [32, 27], (2.1) contains an additional matrix **P**, allowing for a general unknown correspondence between the samples in **X** and **Y**, which is the case in most of our applications. To solve for (2.1), SMAI-align adopted an iterative spectral algorithm that alternatively solves for **P** and (*β, γ*, **R**) using high-dimensional Procrustes analysis. The final solution 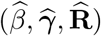 then gives a good similarity transformation aligning the two datasets in the original feature space. In particular, to improve robustness and reduce the effects of potential outliers in the data on the final alignment function, in each iteration we remove some leading outliers from both datasets, whose distances to the other dataset remain large. Moreover, to allow for integration of datasets containing partially shared structures (up to a user-specified threshold, see Methods), users may also request SMAI-align to infer the final alignment function only based on the identified maximal correspondence subsets, rather than the whole datasets. This makes the alignment more robust to local structural heterogeneity. SMAI-align returns an integrated dataset containing all the samples, along with the similarity transformation, which are interpretable and readily used for various downstream analyses (Figure 1b). The idea of SMAI-align is closely related to that of SMAI-test: the passing of SMAI-test essentially renders the goodness-of-fit of the model underlying SMAI-align algorithm, and therefore ensures its performance. In addition, since SMAI-align essentially learns some underlying similarity transformation, based on which all the samples are aligned, the algorithm is easily scalable to very large datasets. For example, one can first infer the alignment function by applying SMAI-align to some representative subsets of the datasets, and then use it to align all the samples.

To empirically evaluate the statistical validity of SMAI-test and the consistency of SMAI-align, we generate simulated data based on some signal-plus-noise matrix model with various signal structures, batch effects, and sample sizes (Supplementary Notes). Our simulation results indicate that SMAI-test has desirable type I errors across all the simulation settings, that is, achieving the nominal low probability (0.05) of rejecting the null hypothesis when the datasets are truly alignable (Table S1, Methods). We also evaluate the performance of SMAI-align in recovering the true alignment function by measuring the estimation errors for each of the true parameters (*β*^*∗*^, ***γ***^*∗*^, **R**^*∗*^) generating the data (Methods). We find that increasing the sample sizes leads to reduced estimation errors in general (Figure S2b), suggesting statistical consistency of SMAI-align.

### SMAI robustly determines alignability between diverse single-cell data

We apply SMAI-test to diverse single-cell data integration tasks and demonstrate its robust performance in determining the alignability between the datasets. The detailed information about each dataset and the corresponding test results are summarized in Table 1. In particular, our real and synthetic datasets involve diverse tissues including human livers, human pancreas, human blood (peripheral blood mononuclear cell, PBMC), human lung, human mesenteric lymph nodes (MLN), mouse brain, mouse primary visual cortex (VISP), and mouse gastrulation, and contain multiple modalities measured by various sequencing technologies, such as single-cell transcriptomics (10X Genomics, Smart-seq, Smart-seq2, Drop-seq, and CEL-seq2), spatial transcriptomics (seqFISH, ISS and ExSeq), and chromatin accessibility (ATAC-seq and 10X Genomics). The 11 integration tasks cover six different scenarios that arise commonly from single-cell research, including (1) integration across different samples with the same cell types, (2) integration across different samples with partially overlapping cell types, (3) integration across samples with non-overlapping cell types, (4) integration across different sequencing technologies, (5) integration across different tissues, and (6) integration of single-cell RNA-seq and spatial-transcriptomic data. For each task, we test for partial alignability between each pair of datasets, determining whether at least 60% of the cells are alignable in the sense of our null model (Method). Among them, three out of eleven integration tasks, with zero or very low proportions (≤ 37%) of cells under the overlapping cell types, are taken as negative controls (Tasks Neg1-Neg3), whose alignability is doubtful in general; the rest of the tasks, including five non-spatial integration tasks (Tasks Pos1-Pos5) and three spatial integration tasks (Tasks PosS1-PosS3), are taken as positive controls, whose alignability is expected due to the association with the same tissue and/or largely overlapping cell types.

**Table 1:** Summary of single-cell datasets analyzed in this paper. The p-values indicate statistical significance against the null hypothesis that the two datasets are at least partially (60%) alignable.

| task id | # of cells | # of genes | same technology (Y-yes; N-no) | reference |
| --- | --- | --- | --- | --- |
| Neg1 | (1272, 1015) | (34363, 34363) | synthetic | [71] |
| Neg2 | (904, 1226) | (15148, 15148) | Y(10X Genomics) | [57] |
| Neg3 | (4000, 4000) | (36601, 36601) | Y(10X Genomics) | [25] |
| Pos1 | (2364, 2244) | (34363, 34363) | N(Smart-seq2, CEL-Seq2) | [71] |
| Pos2 | (2035, 2134) | (34363, 34363) | synthetic | [71] |
| Pos3 | (3362, 3222) | (33694, 33694) | N(10X Genomics v2, v3) | [24] |
| Pos4 | (3618, 3750) | (3429, 3429) | Y(ATAC-seq) | [57] |
| Pos5 | (2406, 2410) | (15148, 15148) | Y(10X Genomics) | [57] |
| PosS1 | (14891, 4651) | (36, 29452) | N(seqFISH, 10X Genomics) | [55, 33] |
| PosS2 | (6000, 34041) | (119, 34041) | N(ISS, Smart-seq) | [34, 80, 56] |
| PosS3 | (1154, 34041) | (42, 34041) | N(ExSeq, Smart-seq) | [4, 80, 56] |

| task id | same tissue (Y-yes; N-no) | % of samples under overlapping cell types | p-values |
| --- | --- | --- | --- |
| Neg1 | synthetic | 0% | $< 10^{-6}$ |
| Neg2 | Y(human lung) | 26% | $< 10^{-3}$ |
| Neg3 | N(human liver, human MLN) | 37% | $< 10^{-9}$ |
| Pos1 | Y(human pancreas) | 100% | 0.44 |
| Pos2 | synthetic | 84% | 0.50 |
| Pos3 | Y(human PBMC) | 99% | 0.78 |
| Pos4 | Y(mouse brain) | 100% | 0.88 |
| Pos5 | Y(human lung) | 100% | 0.14 |
| PosS1 | Y(mouse gastrulation) | — | 0.84 |
| PosS2 | Y(mouse VISP) | — | 0.18 |
| PosS3 | Y(mouse VISP) | — | 0.38 |

SMAI-test assigns significant p-values (i.e., p-value < 0.01) to all the negative control, correctly detecting their unalignability against our null model. The positive control tasks are assigned non-significant p-values, passing its (partial) alignability test as expected. For the five non-spatial integration tasks (Tasks Pos1-Pos5), SMAI-test confirms the alignability between datasets from the same tissues and largely overlapping (≥ 84%) cell types but possibly different sequencing technologies. For the three spatial tasks (Tasks PosS1-PosS3), SMAI-test confirms the alignabiliy of the paired single-cell RNA-seq and spatial-transcriptomic data from the same tissue, justifying its wide use for downstream analyses such as prediction of unmeasured spatial genes (Fig 5).

### Necessity of certifying data alignability prior to integration

In the absence of a principled procedure for determining the alignability between datasets, the existing integration methods often end up forcing alignment between any datasets by significantly distorting and twisting each original dataset (Figure 2). Specifically, for each of the negative control tasks Neg1-Neg3, we apply five popular existing integration methods (Scanorama, Harmony, LIGER, fastMNN, and Seurat) to obtain the integrated datasets, and then evaluate how well the relative distances between the cells within each dataset before integration are preserved in the integrated datasets. As a result, we find overall low correlations (Kendall’s tau correlation on average 0.64 across the five methods and three negative control tasks Neg1-Neg3, as compared with 0.9 achieved by SMAI-align across Pos1-Pos5 on average) between the pairwise distances with respect to each cell before and after data integration (Figure 2b). In addition, we observe many cases of false alignment of distinct cell types from different datasets, and sometimes serious distortion and dissolution of individual cell clusters under the same cell type. For example, in Task Neg1, we find the false alignment of ductal cells and acinar cells by Scanorama, Harmony, and fastMNN, the alignment of alpha cells and gamma cells by Scanorama, fastMNN and Seurat, and the alignment of alpha cells and beta cells by Seurat (Figure 2a); we also observe significant distortion or dissolution of the alpha cell cluster and the beta cell cluster after integration by Harmony, LIGER, and fastMNN, as compared with the original datasets (Figure 2a). As such, the final integration results can be highly problematic and unfaithful to the original datasets, which may lead to erroneous conclusions from downstream analysis. SMAI-test is able to detect the lack of alignability (i.e., significant p-values) between these datasets, alerting users that data integration is not reliable.

**Figure 2.**
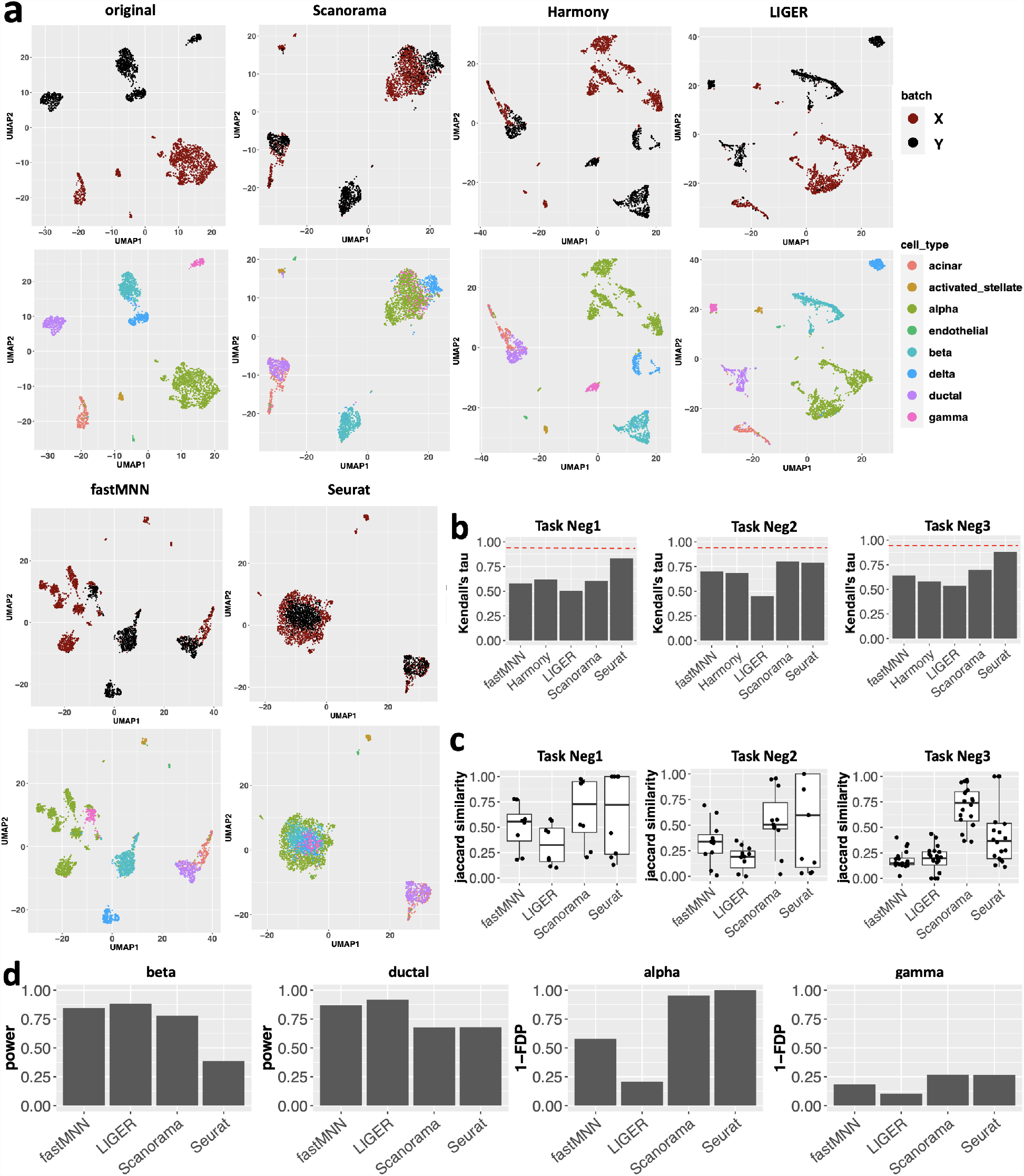
Enforcing uncertified data integration may cause false alignment, serious distortions and misleading inferences. UMAP visualizations of the original (pooled) data under negative control Task Neg1, and the integrated data as obtained by five popular methods (Scanorama, Harmony, LIGER, fastMNN and Seurat). For each method, the top figure is colored to indicate the distinct datasets being aligned, whereas the bottom figure to indicate different cell types. Similar plots for Neg2 and Neg3 are shown in Figure S3. (b) Under the three negative control tasks, we show bar plots of average Kendall’s tau correlations between pairwise distances with respect to each sample in both datasets before and after data integration, as achieved by each methods. The red dashed line benchmarks the average Kendall’s tau correlation of 0.9 achieved by SMAI-align over the positive control tasks Pos1-Pos5. (c) Boxplots of Jaccard similarity between the set of differentially expressed (DE) genes associated to a distinct cell type detected based on the integrated data and the DE genes based on the original data. Each point represents a distinct cell type. (d) For Task Neg1, we show some representative bar plots of (1*−*false discovery proportion) (1*−*FDP) and the power of detecting DE genes for some cell types based on the integrated data. Harmony is not included in (c) and (d) as its integration is only achieved in the low-dimensional space. Notably, SMAI-test correctly detects that all the datasets in Task Neg1-Neg3 are not alignable.

To evaluate the possible effects on downstream analysis, we focus on one important application following data integration, that is, the identification of differentially expressed (DE) genes for each cell type. We consider the above four integration methods (Harmony is not included here as it only produces integrated data in the low-dimensional space). For the three negative control tasks, we find that for many cell types, the set of DE genes identified based on the integrated data can have little overlap with those identified based on the original datasets (Figure 2c and Figure S4). These discrepancies are likely artifacts caused by the respective integration methods. For instance, in Task Neg1, we find that, compared with other methods, the integrated data based on Seurat has lower power in detecting DE genes for beta cells and ductal cells, which may be a result of more serious entanglement of beta and ductal cells with other cell types as created by Seurat integration (Figure 2d). Similarly, we find that the integrated datasets by fastMNN and LIGER data have higher false discovery proportions (FDPs) in detecting DE genes for alpha cells compared with other methods, which is likely a consequence of the distorted or broken alpha cell clusters after alignment (Figure 2d). In this regard, SMAI-test may be used prior to any integration task, to recognize any intrinsic discrepancy between the datasets, and avoid potentially misleading inferences and conclusions.

### SMAI enables principled structure-preserving integration of single-cell data

For the five non-spatial positive control tasks (Tasks Pos1-Pos5), whose datasets are rendered as alignable by SMAI-test, we further apply SMAI-align to obtain the integrated datasets (Figure 3a) and compare the quality of alignment with the above five existing methods. We find SMAI-align has uniformly better performance in preserving the within-data structures after integration (i.e., highest correlations between the pairwise distances with respect to each sample before and after integration), while achieving overall the best performance in removing the unwanted variations between the datasets (i.e., higher similarity in expression profiles for the same cell types across batches) (Figure 3a). In particular, these features are consistent across multiple evaluation metrics, including Kendall’s tau correlation and Spearman’s rho correlation for structure preservation, and the Davies-Bouldin (D-B) index [23, 18, 91, 36, 59] and the inverse Calinski-Harabasz (C-H) index [15, 54, 43, 19, 41] for batch effect removal (Figure S5). The advantages of these indices over other metrics such as LISI or ARI are explained in Methods section. The desirable alignment achieved by SMAI and its advantages over the existing methods is further supported by visualizing low-dimensional embeddings of the integrated data. In Figure 3b, we observe that across all five tasks, SMAI-align in general achieves good alignment of cells from different datasets under the same cell types. In contrast, distortions and misalignment of certain cell types are found for existing methods. For instance, in Task Pos2, we observe false integration of gamma cells and beta cells by Harmony and Seurat, and significant distortion, that is, stretching and creation of multiple artificial subclusters, of the alpha cell cluster by LIGER and fastMNN (Figure S6a). As another example, for Task Pos4, we observe strong distortion and artificial clustering of the excitatory neurons and the inhibitory neurons, in the data output by Harmony, LIGER, and fastMNN (Figure S6b). In general, compared with the existing methods, the integrated datasets obtained by SMAI-align are overall of higher integration quality, and less susceptible to technical artifacts, structural distortions, and information loss, making them more reliable for downstream analyses.

**Figure 3.**
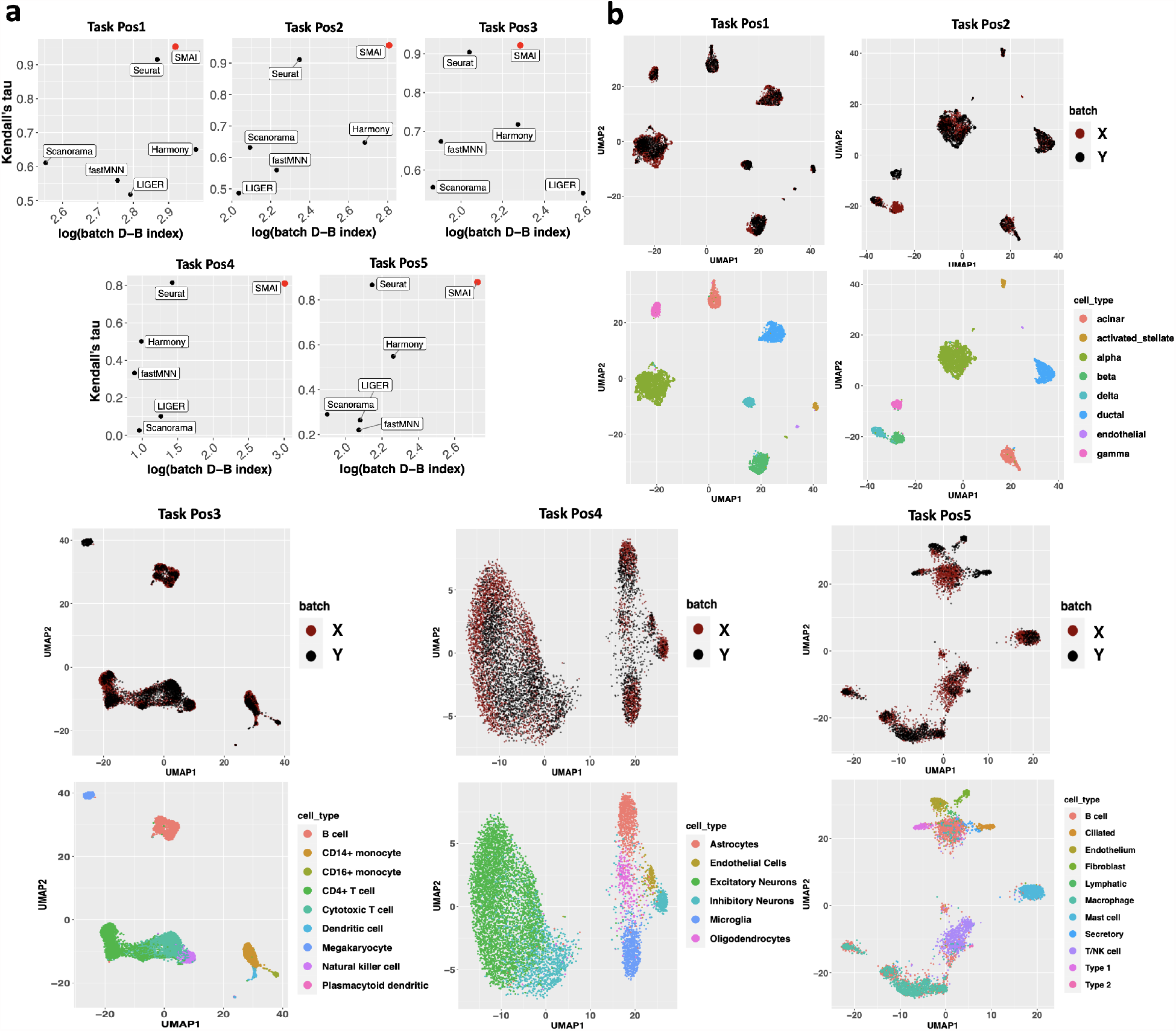
Performance of SMAI-align on the five positive control integration tasks. (a) Compared with five existing algorithms (black), SMAI-align (red) has uniformly better performance in preserving the within-data structures after integration while achieving overall the best performance in removing unwanted variations. The former is characterized by the highest Kendall’s tau correlations between the relative distances with respect to each sample before and after integration (y-axis), whereas the latter is reflected by higher values of the batch-associated Davies-Bouldin (D-B) index (x-axis shown in log-scale). See Figure S5 for similar results under other metrics. (b) UMAP visualizations of the integrated data as obtained by SMAI-align. For each integration task, the top figure is colored to indicate the datasets being aligned, whereas the bottom figure is colored to indicate different cell types. See also Figure S6 for UMAP visualizations associated with other integration methods.

### SMAI improves reliability and power of differential expression analysis

A common and important downstream analysis following data integration is to identify the marker genes associated with individual cell types based on the integrated data. To demonstrate the advantage of SMAI-align in improving the reliability of downstream differential expression analysis, we focus on the above five positive control tasks and evaluate how many DE genes for each cell type are preserved after integration, and how many new DE genes are introduced after integration. Specifically, for each integrated dataset produced by fastMNN, LIGER, Scanorama, Seurat, or SMAI, we identify the DE genes for each cell type based on the Benjamini-Hochberg (B-H) adjusted p-values, and compare their agreement with the DE genes identified from the individual datasets before integration using the Jaccard similarity index, which accounts for both power and false positive rate in signal detection (Methods). As a result, we find that compared with other methods, SMAI-align oftentimes leads to more consistent and more reliable characterization of DE genes based on the integrated data (Figure 4a).

**Figure 4.**
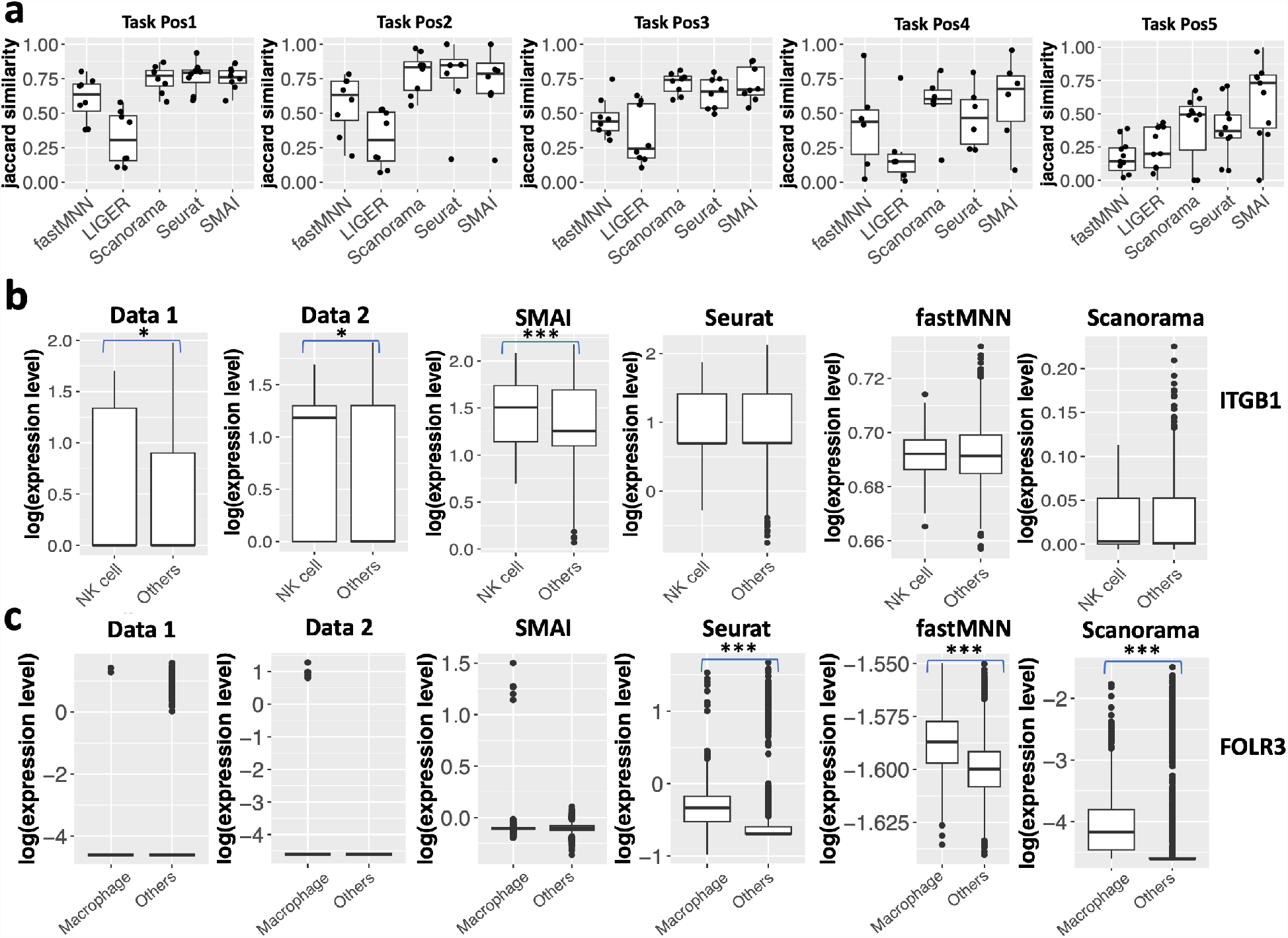
SMAI improves reliability and power of DE analysis. (a) Boxplots of Jaccard similarity between the DE genes for each cell type identified based on the integrated data, obtained by one of the five integration methods, and those identified based on the individual datasets before integration. Each point represents a distinct cell type. The results indicate that SMAI-align oftentimes lead to more consistent and more reliable characterization of DE genes, as compared with other methods. (b) Boxplots of log-expression levels of ITGB1 as grouped by cell types in the two datasets about human PBMCs (Task Pos3: Data 1 contains 219 natural killer (NK) cells and 3143 other cells, and Data 2 contains 194 NK cells and 3028 other cells), and in the integrated datasets (413 NK cells and 6171 other cells) as produced by SMAI-align, Seurat, fastMNN, and Scanorama. The DE pattern of ITGB1 is only preserved by SMAI after integration. (c) Boxplots of log-expression levels of FOLR3 as grouped by cell types in the two datasets about human lung tissues (Task Pos5: Data 1 contains 68 macrophages and 2285 other cells, and Data 2 contains 911 macrophages and 1000 other cells), and in the integrated datasets (979 macrophages and 3285 other cells) as produced by SMAI-align, Seurat, fastMNN, and Scanorama. Artificial DE patterns are created by existing integration methods. The stars above the boxplots indicate statistical significance of DE test. Specifically, * means adjusted p-value *<* 0.05; ** means adjusted p-value *<* 0.01; *** means adjusted p-value *<* 0.001. Harmony and LIGER are not included in (b) and (c) as they do not produce gene-specific integrated data.

Biological insights can be obtained from the improved DE analysis with SMAI-align. For instance, under Task Pos3 concerning human PBMCs, an important protein coding gene ITGB1 (CD29), involved in cell adhesion and recognition [38], has been found differentially expressed in natural killer (NK) cells compared with other cell types in the SMAI-integrated data (adjusted p-value *<* 10^*−*11^), but not in the Seurat-, fastMNN-, or Scanorama-integrated data. In both original datasets before integration, we also find statistical evidence supporting ITGB1 as a DE gene for NK cells (Figure 4b). Interestingly, the functional relevance of ITGB1 to NK cells in determining its cell identity and activation has been reported in a large-scale study using the whole-genome microarray data of the Immunological Genome Project [11]. In this case, the biological signal is blurred and compromised during data alignment by existing methods. See also Figure S7 for more examples. On the other hand, we also find evidence suggesting that SMAI-align is less likely to introduce artificial signals or false discoveries as compared with existing methods. For example, under Task Pos5 concerning human lung tissues, we find almost no expression of gene FOLR3 in macrophages in both datasets before integration. However, after integration, both genes are detected as DE genes for their respective cell types based on the Seurat-, fastMNN-, and Scanorama-integrated datasets, but not based on the SMAI-integrated dataset (Figure 4c). Similar examples are shown in Figure S8. Such artificial signals are likely consequences of overly distorted expression profiles of these cell types created by existing integration methods that locally search for the best alignment. Such a limitation can be overcome by SMAI-align, which is a global alignment method.

To strengthen our argument on the advantage of SMAI in improving downstream DE analysis, we also carry out simulation studies by generating pairs of datasets, each containing 2000 cells of 12 different cell types (clusters) and expression levels of 1000 genes, with different cell type proportions. We create batch effects between the two datasets mimicking those observed in real datasets (Methods). For two different simulation settings, we evaluate the Jaccard index, the false discovery rate, and the power, of the subsequent DE analysis with respect to the true marker genes, based on each of the integrated datasets. This analysis further confirms the advantage of SMAI-align over the other four alternative methods in precisely characterizing the true marker genes, as measured by the above three metrics (Figure S9). In particular, our simulation study demonstrates the tendency of creating more false positives by the existing methods as compared with SMAI-align.

### SMAI improves integration of scRNA-seq with spatial transcriptomics

Another important application of the data integration technique is the imputation of the spatial expression levels of unmeasured transcripts in single-cell spatial transcriptomic data. Spatial transcriptomics technologies extend high-throughput characterization of gene expression to the spatial dimension and have been used to characterize the spatial distribution of cell types and transcripts across multiple tissues and organisms [5, 62, 39, 60, 12, 63, 86]. A major trade-off across all spatial transcriptomics technologies is between the number of genes profiled and the spatial resolution such that most spatial transcriptomics technologies with single-cell resolution are limited to the measurement of a few hundred genes rather than the whole transcriptome [52]. Given the resource-intensive nature of single-cell spatial transcriptomics data acquisition, computational methods for upscaling the number of genes and/or predicting the expression of additional genes of interest have been developed, which oftentimes make use of some paired single-cell RNA-seq data. Among the existing prediction methods, an important class of methods [1, 74, 3, 87] are based on first aligning the spatial and RNA-seq datasets and then predicting expression of new spatial genes by aggregating the nearest neighboring cells in the RNA-seq data. Applications of these methods have been found, for example, in the characterization of spatial differences in the aging of mouse neural and glial cell populations [3], and recovery of immune signatures in primary tumor samples [84]. As a key step within these prediction methods, we show that data alignment achieved by SMAI may lead to improved performance in predicting unmeasured spatial genes. To ensure fairness, we compare various prediction workflows which only differ in the data integration step (Methods). For the three spatial positive control tasks (PosS1-PosS3), we withhold each gene from the spatial transcriptomic data, and compare its actual expression levels with the predicted values based on the aforementioned two-step procedure where the data alignment is achieved by one of the six methods (LIGER, Scanorama, Harmony, Seurat, fastMNN, and SMAI-align). Due to the intrinsic difficulty of predicting some spatial genes that are nearly independent of any other genes, we only focus on predicting the first half of spatial genes that have higher correlations with some other genes. Our analysis of the three pairs of datasets yields the overall best predictive performance of the SMAI-based prediction method (Figure 5).

### SMAI’s interpretability reveals insights into the sources of batch effects

Unlike the existing methods, SMAI-align not only returns the aligned datasets, but also outputs explicitly the underlying alignment function achieving such alignment, which enables further inspection and a deeper understanding of possible sources of batch effects. Specifically, recall that the final alignment function obtained by SMAI-align consists of a scaling factor 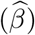, a global mean-shift vector 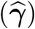, and a rotation matrix 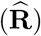; each of them may contain important information as to the nature of the corrected batch effects. For example, applying SMAI-align to the human pancreas data (Task Pos1) leads to an alignment function in terms of 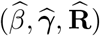, linking the CEL-Seq2 dataset to the Smart-Seq2 dataset. The scaling factor 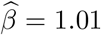 suggests little scaling difference between the two datasets. However, the obtained global mean-shift vector 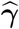 highlighted a sparse set of genes such as SST, CTSD, PEMT, and REG1A, affected by the batch effects (Figure 6a). In particular, for both datasets, we observe similar patterns in the relative abundances of the transcript across different cell types (Figure 6b), suggesting the batch effects on these genes are relatively uniform across cell types. Moreover, while SST, CTSD, PEMT, and REG1A are all DE genes associated with some cell types, SMAI-align doesn’t affect DE results after integration due to its ability to distinguish and remove such global discrepancy. Similar observations can be made on other integration tasks such as human PBMCs (Task Pos3, Figure S10). As for the rotation matrix 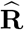, it essentially captures and remove the batch effects altering the gene correlation structures in each dataset (Figure S11). The obtained SMAI-align parameters 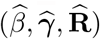 can also be converted into distance metrics to quantify and compare the geometrically constitutive features of the batch effects (Figure S12a) and their overall magnitudes (Figure S12b).

**Figure 5.**
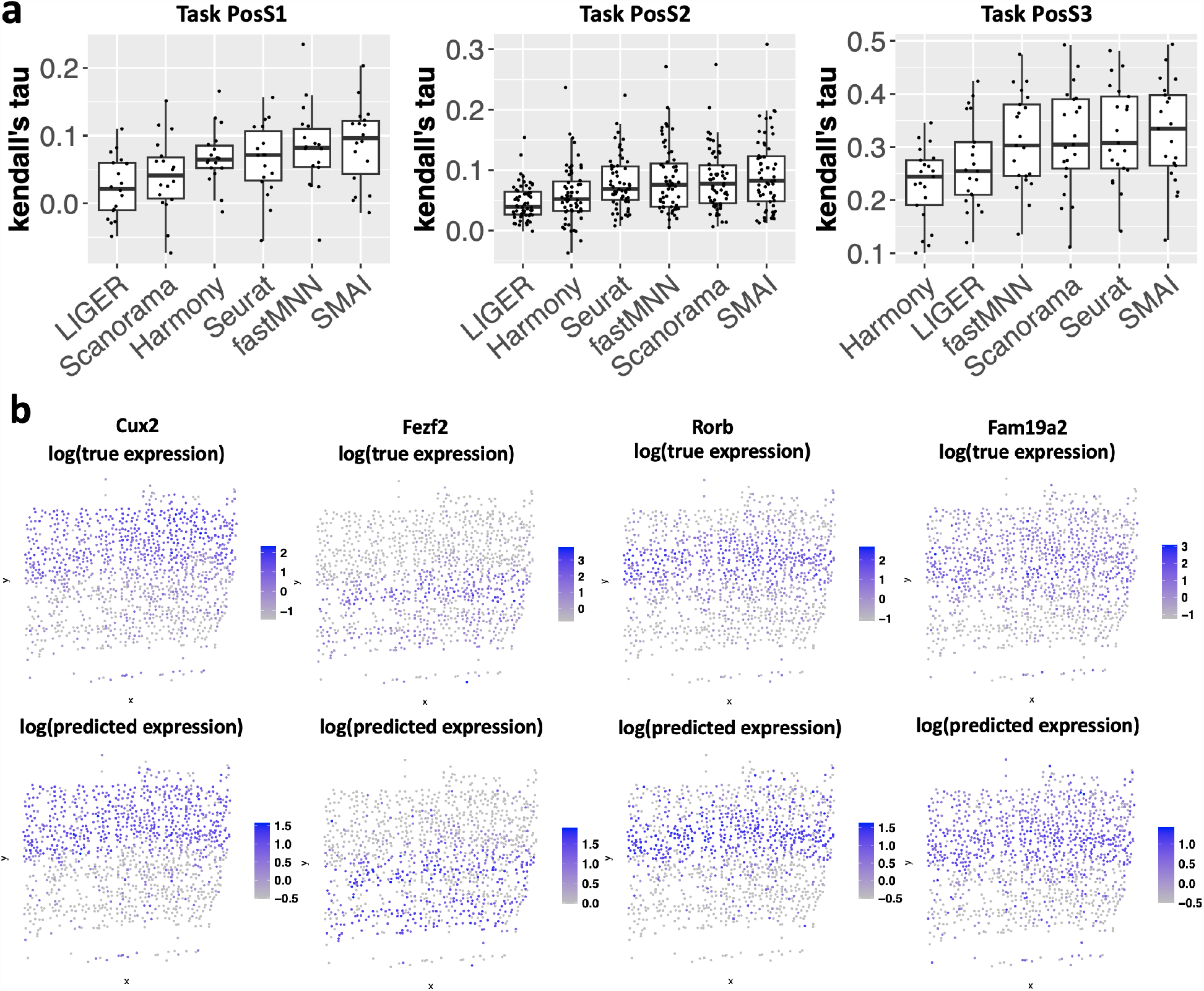
SMAI improves prediction of single-cell spatial transcriptomic data. (a) Boxplots of kendall’s tau correlation between the actual expression levels of the spatial genes and their predicted values based on the two-step procedure (alignment followed by *k* nearest neighbor regression) where the data alignment is achieved by LIGER, Scanorama, Harmony, Seurat, fastMNN, or SMAI-align. Each point represent a distinct spatial gene. The methods are ordered according to their median predictive performance, showing the overall best performance of SMAI. (b) Examples of true expression levels some spatial genes from Task PosS3, presented according to the cells’ spatial layout, and their predicted values based on SMAI-align. The colors are in log-scale.

**Figure 6.**
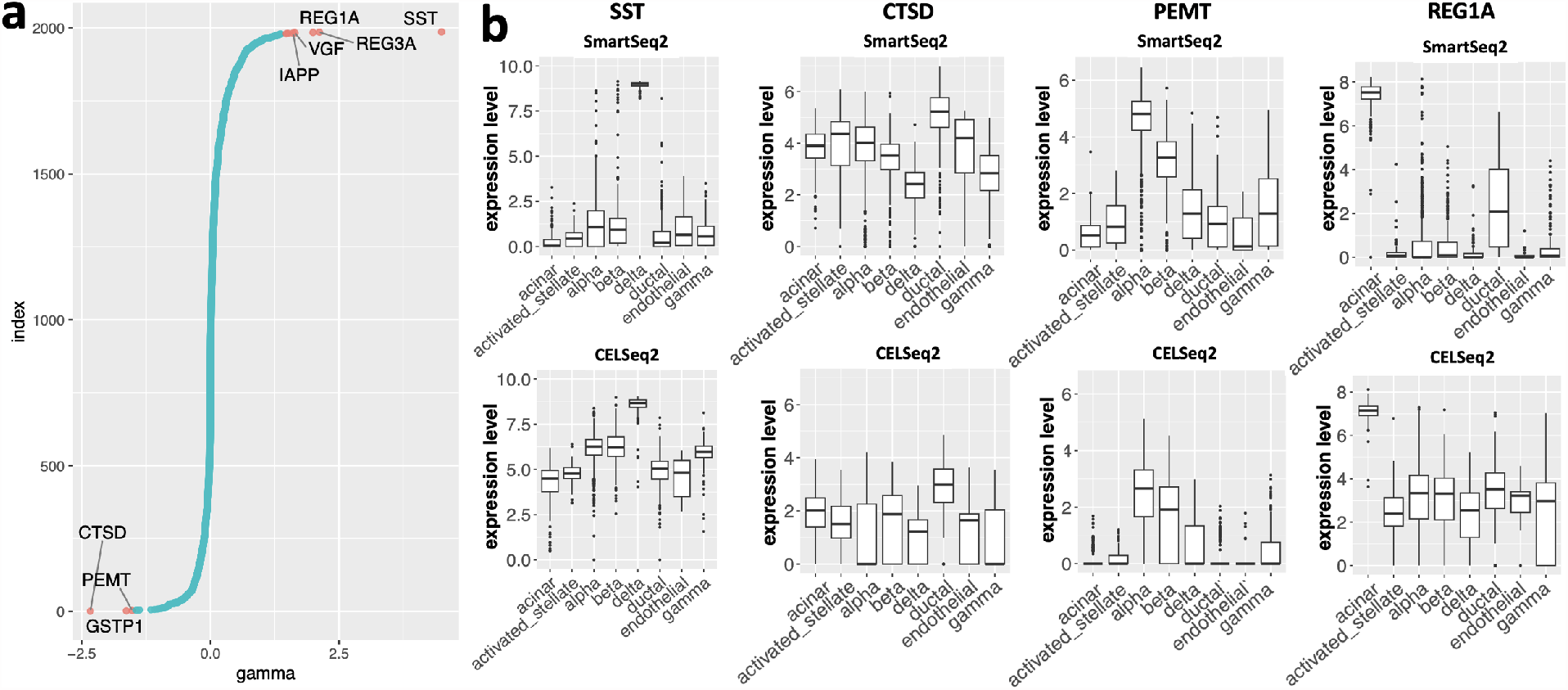
SMAI’s interpretability gives insights into the batch effects. (a) Visualization of the estimated mean-shift vector 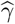 from integrating the pancreatic data (Task Pos1), whose components are ordered from the smallest (bottom) to the largest (top). In 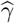 only a sparse set of genes such as SST, CTSD, PEMT, and REG1A are affected by the batch effects. (b) Boxplots of expression levels of some genes highlighted in (a), as group by different cell types. Top: Smart-Seq2 dataset with *n* = 2364. Bottom: CEL-Seq2 dataset with *n* = 2244. Note that SST, CTSD, PEMT, and REG1A are all DE genes. The batch effects on these genes are relatively uniform across cell types, and therefore SMAI-align doesn’t affect DE results after integration.

## 3 Discussion

In this study, we develop a spectral manifold alignment and inference algorithm that can determine general alignability between datasets and achieve robust and reliable alignment of single-cell data. The method explores the best similarity transformation that minimizes the discrepancy between the datasets, and thus is interpretable and able to reveal the source of batch effects. In terms of computational time, standard SMAI requires a similar running time as existing methods such as Seurat (Methods and Figure S13a). A key hyperparameter of SMAI is the number of informative eigenvalues *r*_max_ associated with each data matrix, which can be determined consistently using data-driven procedures incorporated in our algorithm (Methods). Other hyperparameters such as the number of iterations and the number of outliers to be removed per iteration, have recommended default values (Methods), shown to work robustly in our benchmark datasets. Moreover, the availability of a closed-form alignment function allows for fast alignment of very large datasets, by first learning the alignment function from some representative subsets and then apply it to the whole dataset.

SMAI has a few limitations that deserve further development. First, although our restriction to the similarity class already yields competent performance over diverse applications, extending to and allowing for more flexible nonlinear transformations may lead to further improvement, especially in tracking and addressing local discrepancies associated with particular cell types. Second, the current method only makes use of the overlapping features in both datasets. However, in many other applications, such as integrative analysis of single-cell DNA copy number variation data and single-cell RNA-seq data [92, 90], the features are related but not shared in general. Nevertheless, the unique features in each dataset may still be useful for integrative multiomic analysis, or data alignment in some different sense. Third, although the current alignment framework mainly concerns testing and aligning two datasets, direct extension to multiple datasets is available, which is achieved by applying SMAI in a sequential manner, based on some pre-specified order for integrating the datasets. However, we also point out that given the nature of our alignability test and the spectral alignment algorithm, extension to the simultaneous (non-sequential) testing and alignment of multiple datasets may be achieved by replacing the Procrustes analysis objective in (2.1) with a generalized Procrustes analysis objective involving all the available datasets and multiple alignment functions [32, 27]. Expanding SMAI’s scope of applications along these directions are interesting directions of future work.

## 4 Methods

### Basic SMAI algorithm

For clarity and to ease our presentation, we first introduce the basic version of SMAI algorithms (i.e., basic SMAI-test and basic SMAI-align) along with their theoretical properties. Specifically, we first consider testing for complete alignability and obtaining the full alignment of a pair of datasets with matched sample size. Extensions to datasets with general unequal sample size is straightforward and will be discussed subsequently. In later sections, we show that the complete version of SMAI, which flexibly allows for partial alignment, as used in our data analysis, can be obtained with slight modifications of these basic algorithms.

### SMAI-test algorithm

The basic SMAI-test algorithm requires as input the data matrices **X** ∈ ℝ^*d×n*^ and **Y** ∈ ℝ^*d×n*^, each containing *n* cells and *d* genes, a pre-determined parameter *r*_max_ corresponding to the number of leading eigenvalues to be used in the subsequent test, and a global rescaling factor *b >* 0. In principle, the number *r*_max_ should reflect the dimension of the underlying true signal structure of interest, which is unknown in most applications. Nevertheless, there are several theoretically justified approaches available to determine *r*_max_, as we will discuss below. The global scaling factor *b >* 0 adjusts for the potential scaling difference between the two matrices, which can be estimated by, for example,

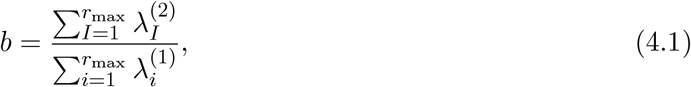

where 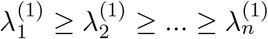 and 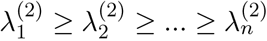 are the ordered eigenvalues of the sample covariance matrices of **X** and **Y**, respectively. The formal procedure of the test is summarized in Algorithm 1, which produces a p-value signifying the strength of statistical evidence against the null hypothesis about the overall alignability of the two datasets. The rigorous statement of the null hypothesis, the underlying statistical model, as well as the theoretical guarantee of the proposed test, will be presented in the next subsection.

#### Algorithm 1 Basic SMAI-test

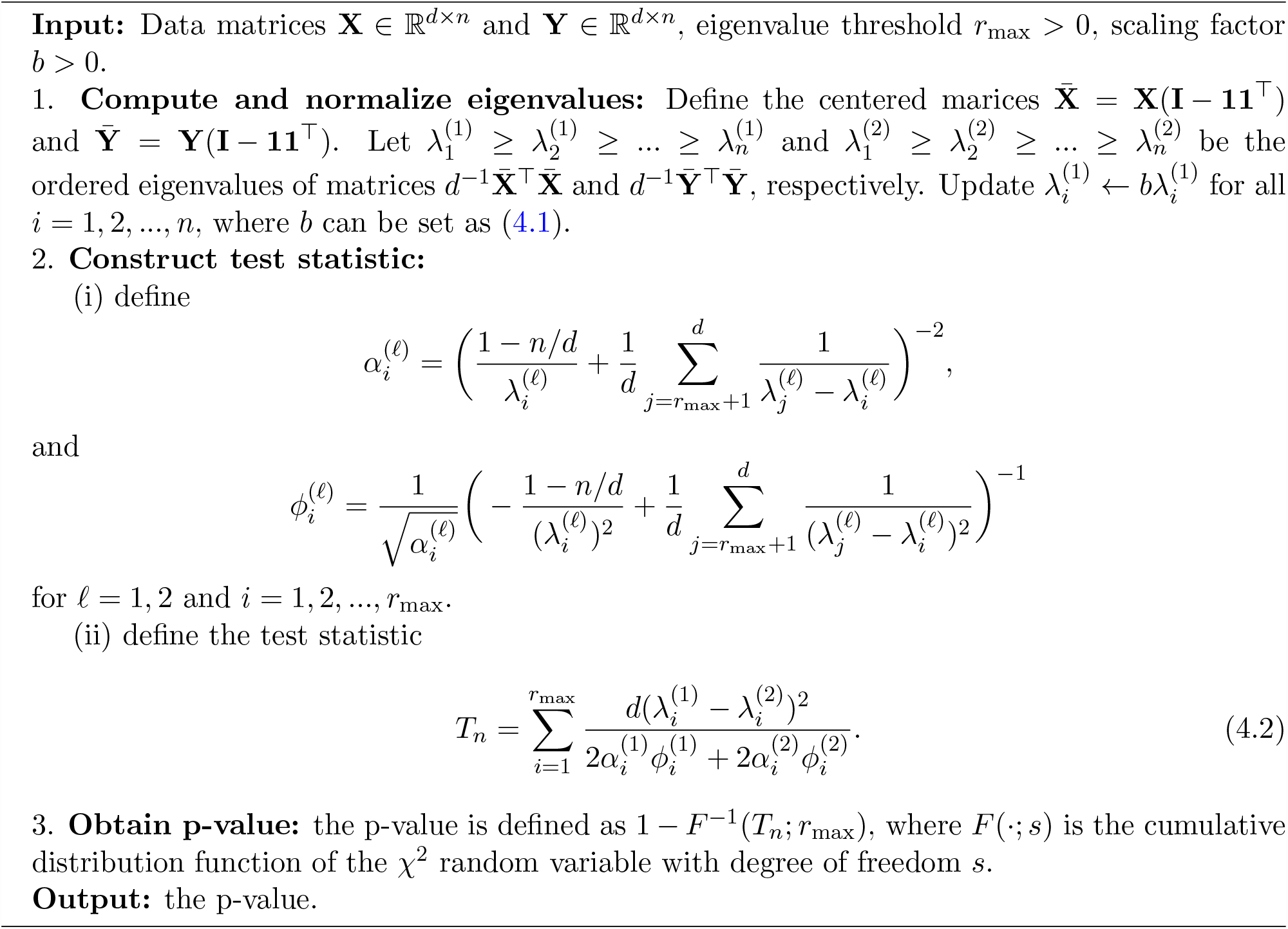

### Theoretical guarantee of SMAI-test

We first introduce our assumption on the centered matrices 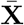 and 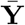 in Algorithm 1, as well as the formal null hypothesis, based on which our testing procedure is developed. Suppose 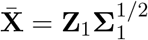 and 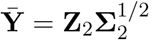, where **Σ**_1_, **Σ**_2_ ∈ ℝ^*n×n*^ are positive definite matrices, and **Z**_1_, **Z**_2_ are independent copies of **Z** = (*z*_*ij*_) ∈ R^*d×n*^ with entries *z*_*ij*_ = *d*^*−*1*/*2^*q*_*ij*_ where the double array {*q*_*ij*_ : *i* = 1, 2, …, *d, j* = 1, 2, …, *n*} consists of independent and identically distributed random variables whose first four moments match those of a standard normal variable. We assume each of **Σ**_1_ and **Σ**_2_ has *r* spiked/outlier eigenvalues, and the remaining bulk eigenvalues have some limiting spectral distribution. In other words, for each **Σ**_*ℓ*_, .*ℓ* = 1, 2, there are exactly *r* eigenvalues 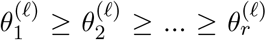 larger than a certain threshold (see (A1)-(A3) in the Supplementary Material for the precise statements), characterizing the dominant global signal structures in the data, whereas for the rest of the eigenvalues 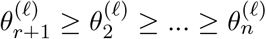 characterizing the remaining signal structures of much smaller magnitude, their empirical distribution (i.e., histogram) has a deterministic limit as *n → ∞*. Moreover, to account for high-dimensionality of the datasets, we assume that the number of genes is comparable to the number of cells, in the sense that *n/d* → *c* ∈ (0, ∞) as *n* → *∞*. The above model is commonly referred as the high-dimensional generalized spiked population model [6, 53, 14, 93], which is widely used to for modeling high-dimensional noisy datasets with certain low-dimensional signal structures. In particular, the key assumption about the spiked eigenvalue structure is supported by empirical evidences from real-world single-cell datasets (Figure S1 and references [50, 49]). Essentially, such a model ensures the existence and statistical regularity of some underlying low-dimensional signal structure. For instance, it implies that when the signal strength of the low-dimensional structure is strong enough, that is, when top eigenvalues 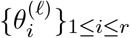 are large enough, the underlying low-dimensional signal structure would be roughly captured by the leading *r* eigenvalues and eigenvectors of 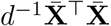 and 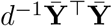. Unlike many earlier work [40, 7, 67, 8], where the bulk eigenvalues 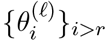 are assumed to be identical or simply ones, the current framework allows for more flexibility as to the possible heterogeneity in the signal and/or noise structures.

Suppose the eigendecompositions of **Σ**_1_ and **Σ**_2_ are expressed as

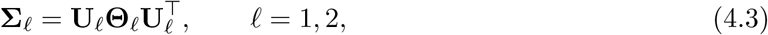

where 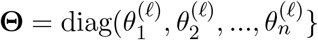 is a diagonal matrix containing the eigenvalues, and the columns of 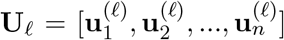 are the corresponding eigenvectors. By definition, the (centered) low-dimensional structure associated with 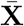 or 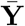 can be represented by the leading *r* eigenvectors of **Σ**_*ℓ*_ weighted by the square root of their corresponding eigenvalues, that is,

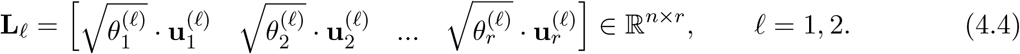

With these, the null hypothesis about the general alignability between **X** and **Y** can be formulated as the alignability between their low-dimensional structures **L**_1_ and **L**_2_ up to a possible rescaling and a rotation (note that translation is not needed as **L**_*ℓ*_’s are already centered). Formally, the null hypothesis under the above statistical model states that

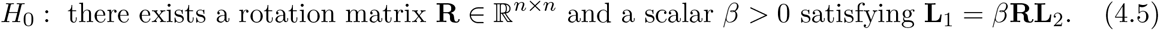

To develop a statistical test against such a null hypothesis, we notice that under *H*_0_, it necessarily follows that 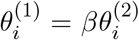 for all 1 *≤ i ≤ r*. As a result, it suffices to develop a statistical test that can evaluate if the spiked eigenvalues of **Σ**_1_ and **Σ**_2_ are identical up to a global scaling factor. This leads us to the proposed test in Algorithm 1. The following theorem, whose proof can be found in the Supplementary File, theoretically justifies the proposed test and ensures its statistical validity in terms of type I errors.

#### Theorem 4.1.

*Suppose* 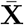 *and* 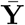 *are independent and satisfy the above high-dimensional generalized spiked population model. Let* 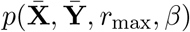 *be the p-value returned by Algorithm 1 with* (*r*_max_, *b*) = (*r*, 1*/β*). *Under the null hypothesis H*_0_ *in (4*.*5), for any* 0 *< α <* 1*/*2, *it holds that*

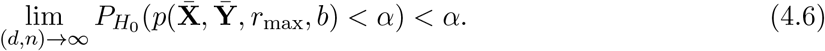

### Advantages of SMAI-test as suggested by statistical theory

A few remarks are in order concerning the advantages of SMAI-test implied by our theory. Firstly, unlike many classical statistical tests for shape conformity, in which *d* is assumed to be fixed (effectively a small integer) as *n → ∞*, the statistical validity of the proposed test is guaranteed (Theorem 4.1) under the so-called high-dimensional asymptotic regime, where *d* is comparable to *n* as *n → ∞*. This makes our testing framework more useful and coherent with the commonly high-dimensional single-cell genomic data. Secondly, SMAI-test is developed based on the spiked population model, which essentially assumes a few large (“spiked”) eigenvalues encoding the dominating low-dimensional signal structure, followed by a large number of smaller bulk eigenvalues. Such a spectral property has been widely observed in single-cell genomic data, and allows for straightforward empirical verification (Figure S1). Thirdly, in contrast with many existing tests whose validity relies on rather rigid signal/noise regularity (e.g., identical bulk eigenvalues), the proposed test incorporates a self-calibration scheme in constructing the test statistic *T*_*n*_, allowing for general bulk eigenvalues and is thus applicable to a wider range of real-world settings. Finally, the proposed alignability test is data-centric and does not require side information such as cell type, tissues, or species identities. This ensures an unbiased characterization of the similarity between datasets according to their actual geometric measures rather than potentially problematic annotations.

### SMAI-align algorithm

The basic SMAI-align method is summarized in Algorithm 2. The algorithm requires as input the normalized data matrices **X** and **Y**, the eigenvalue threshold *r*_max_, the maximum number of iterations *T*, and the outlier control parameter *k*. The algorithm starts with a denoising procedure during which the best rank *r*_max_ approximations (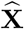 and 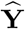) of the data matrices are obtained. Like the determination of *r*_max_, there are multiple ways available to denoise a high-dimensional data matrix with low-rank signal structures [64, 31, 51]. To ensure both computational efficiency and theoretical guarantee, we adopt the hard-thresholding denoiser, summarized in Algorithm 3. The second step is a robust iterative manifold matching and correspondence algorithm, motivated by the shuffled Procrustes optimization problem (2.1). The use of spectral methods in Steps 2(ii) and 2(iii) generalizes the classical ideas of shape matching and correspondence in computer vision [72, 73, 10] and the theory of Procrustes analysis (see Theorem 4.2 below) to high dimensions. To improve robustness with respect to potential outliers in the data, in each iteration we remove the top *k* outliers from both datasets, whose distances to the other dataset remain large after alignment. *T* and *k* are tunable parameters, about which we find that 3 *≤ T ≤* 5 and 5 *≤ k ≤* 20 work robustly for various datasets analyzed in this paper. In the last step, we determine the direction of alignment by either aligning **Y** to **X**, or aligning **X** to **Y**. In general, such directionality is less important as our similarity transformation is invertible and symmetric with respect to both datasets. As a default setting, our algorithm automatically determines the directionality in Step 3, by pursuing whichever direction that leads to a smaller between-data distance. The software allows the users to specify the preferred directionality, which can be useful in some applications such as in the prediction of unmeasured spatial genes using single-cell RNA-seq data.

#### Algorithm 2 Basic SMAI algorithm

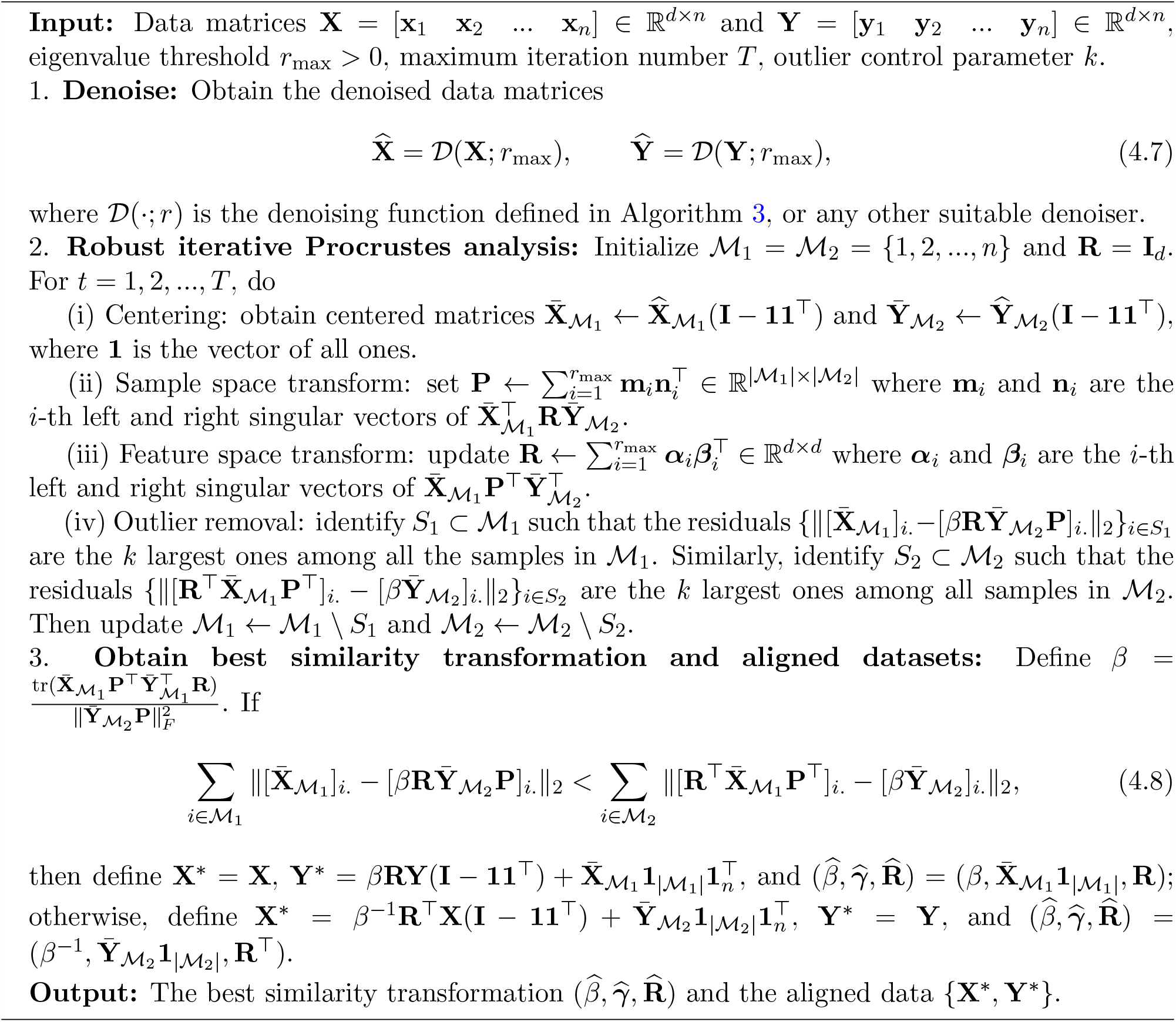

#### Algorithm 3 Hard-thresholding denoiser

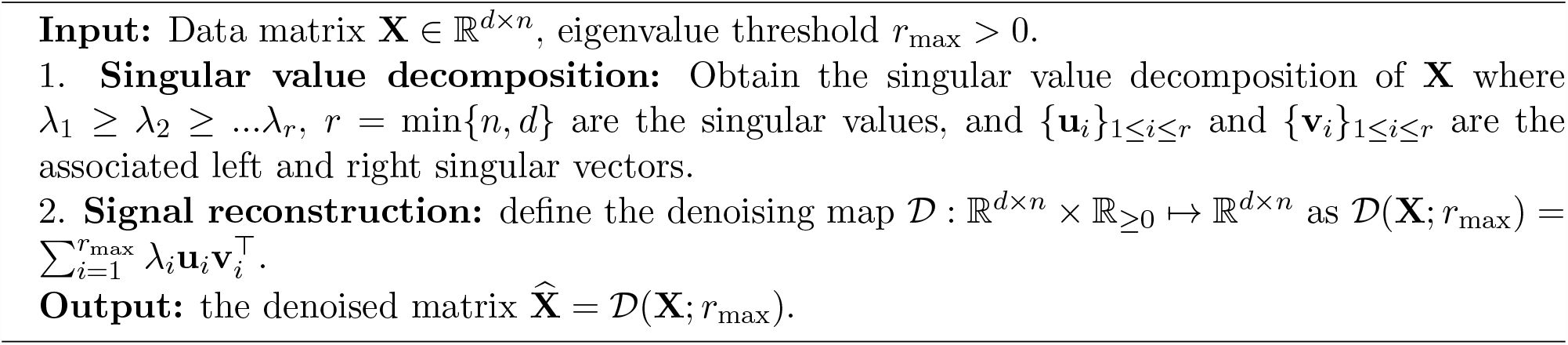

### Rationale and geometric interpretation of SMAI-align

The key step of SMAI-align is an iterative manifold matching and correspondence algorithm. The algorithm alternatively searches for the best basis transformation over the sample space and the feature space. Specifically, in each iteration, the following ordinary Procrustes analyses are considered in order:

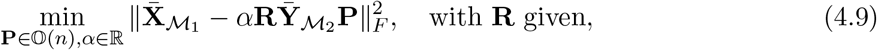

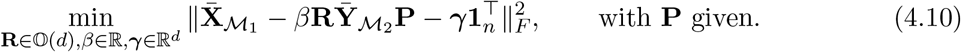

The first optimization problem looks for an orthogonal matrix **P** ∈ 𝕆(*n*) and a scaling factor *α ∈* R so that the data matrix 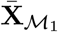 is close to the rescaled data matrix 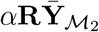, subject to recombination of its samples. In the second optimization problem, a similarity transformation (*β*, ***γ*, R**) is obtained to minimize the discrepancy between the data matrix 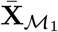 and the sample-matched data matrix 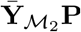. Intuitively, **P** can be considered as a relaxation of permutation matrices allowing for general linear combinations for sample matching between the two data matrices, which may account for differences in sample distributions, such as proportions of different cell types, between the two datasets; **R** represents the rotation needed to align the features between the two matrices. The above optimization problems admit closed-form solutions, which only require a singular value decomposition of some product matrix. The following theorem summarizes some mathematical facts from the classical Procrustes analysis [32, 27].

#### Theorem 4.2.

*Let* **X**_1_, **X**_2_ ∈ ℝ^*p×q*^ *be any matrices, whose column-centered counterparts* 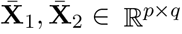 *are such that* 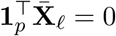 *for* .*ℓ* = 1, 2.

1. *Let* 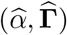 *be the solution to the optimization problem*

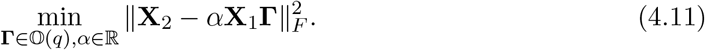

*Then it holds that* 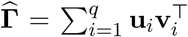, *where* {**u**_*i*_}_1*≤i≤q*_ *and* {**v**_*i*_}_1*≤i≤q*_ *are the ordered left and right singular vectors of* 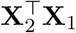, *respectively*.
2. *Let* 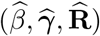 *be the solution to the optimization problem*

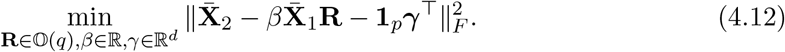

*Then it holds that*

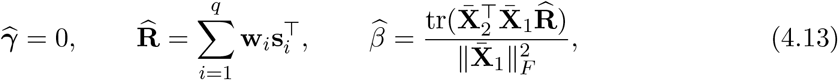

*where* {**w**_*i*_}_1*≤i≤q*_ *and* {**s**_*i*_}_1*≤i≤q*_ *are the ordered left and right singular vectors of* 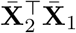, *respectively*.

In particular, Part 1 of Theorem 4.2 explains the rationale behind Steps 2(ii) of Algorithm 2, whereas Part 2 of Theorem 4.2 justifies Step 2(iii), and the form of the estimator for *β* in Step 3 of Algorithm 2. Note that in Algorithm 2 we slightly abuse the notation so that 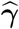 is defined to account for the global mean shift between **X** and **Y**, rather than between 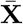 and 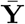 as in Theorem 4.2. In both Steps 2(ii) and 2(iii), to adjust for high-dimensionality and reduce noise, only the leading singular vectors are used to reconstruct the transformation matrices **P** and **R**.

### Determine the rank parameter *r*_max_

In both SMAI-test and SMAI-align, an important parameter to be specified is *r*_max_, the number of leading eigenvalues and eigenvectors expected to capture the underlying signal structures. In the statistical literature, under the spiked population model, various methods have been developed to consistently estimate this value in a data-driven manner [45, 46, 66, 20, 42, 26]. In our software implementation, we provide two options for estimating *r*_max_. The simpler and computationally more effective approach is to consider

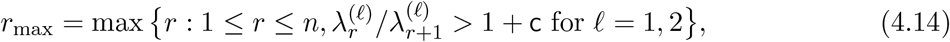

for some small constant c *>* 0 such as 0.01. Theoretical justifications of such a method have been established based on standard concentration argument [89]. A more recent approach, ScreeNOT [26], is also implemented as an alternative; it is theoretically more appealing but requires additional computational efforts.

### Unequal sample sizes

The basci versions of SMAI-test and SMAI-align can be easily adjusted to allow for unequal sample sizes. For example, if 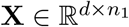 and 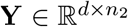 where, say, *n*_1_ *> n*_2_, then one can simply randomly subsample the columns of **X** to obtain 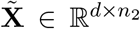 as an input in place of **X**. One advantage of the current framework is the generalizability of the obtained similarity transformation. Specifically, in Algorithm 2, the similarity transform 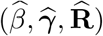, obtained based on subsamples and 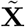 and **Y**, can still be applied to all the samples, leading to the aligned data 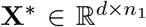 and 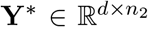 of all the samples. In other words, the subsampling step does not affect the size of the final output of SMAI-align.

### Partial alignability and partial alignment

To allow for more flexibility, a sample splitting procedure is incorporated into SMAI-test, so that one can determine if there exist subsets containing *t*% of the samples in each dataset are alignable up to some similarity transformation. The main idea is to first identify subsets of both datasets, referred to as the maximal correspondence subsets, each containing about *t*% of the samples, expected to be more similar than other subsets, and then perform SMAI-test on these identified subsets, to determine their overall alignability. Specifically, the samples in **X** and **Y** are first split randomly into two parts 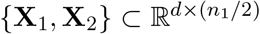 and 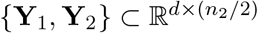, each containing half of the samples. Next, we identify subsets of **X**_1_ and **Y**_1_ containing *t*% of their samples, that are expected to have more similar underlying structures than other possible subsets. This is achieved based on the independent datasets **X**_2_ and **Y**_2_ with the following three steps:

1. Obtain an initial alignment between **X**_2_ and **Y**_2_ using SMAI-align or any existing methods;
2. Identify subsets 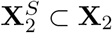 and 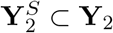, each containing *t*% of the samples, so that 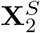 and 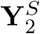 have the most structural similarity; this can be achieved, for example, by searching for the mutual nearest neighbors after alignment; and
3. Identify subsets 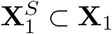 and 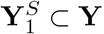, each containing *t*% of the samples, which are nearest neighbors of 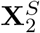 and 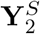, respectively.

With these, the partial alignability between **X** and **Y** can be determined by applying SMAI-test to the submatrices 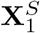 and 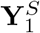. Importantly, by introducing the sample-splitting, the subsequent hypothesis testing is based on the samples independent from those used in the prior subset selection step, making it free from selection bias or double-dipping. As a result, by first conditioning on the subsets {**X**_2_, **Y**_2_} and then integrating out its uncertainty, it can be shown that the statistical validity (controlled type I error) of the final test results still hold as in Theorem 4.1. For practical use, we recommend setting *t* between 50 and 70 to simultaneously retain robustness against local heterogeneity and ensure statistical power with sufficient sample size. Moreover, one should aware that there is a tradeoff between the threshold *t* and the associated p-value: when *t* is below the true proposition of the alignabile samples, the p-values are usually not significant as the null hypothesis about partial alignability remains true; when *t* becomes larger than that, the null hypothesis will be mostly rejected. This relationship can be seen empirically in Figure S13.

In line with the partial alignability test, in our software implementation, we also allow restricting SMAI-align to 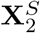 and 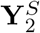 to obtain the inferred similarity transformation. In this way, the final alignment is essentially achieved by focusing on the maximal correspondence subsets rather than the whole set, making the alignment function more robust to local structural heterogeneity.

### Simulation studies: consistency and type I errors

We generate data matrices **X** and **Y** following the simulation setup described in the Supplementary Notes. Under each round of simulation, we apply SMAI-align to obtain estimates of the underlying similarity transformation, as parametrized by (*β*^*∗*^, ***γ***^*∗*^, **R**^*∗*^). The estimated parameters 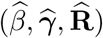 as returned from Algorithm 2, up to possible reversion of the similarity transformation, are evaluated based on the following loss functions

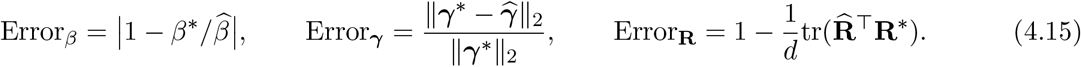

The estimation errors under various samples sizes and signal structures are shown in Figure S2c.

To evaluate the empirical type I error of SMAI-test, we apply SMAI-test to the simulated datasets sharing the same low-dimensional structure subject to some similarity transformation. In each round of simulation, we reject the null hypothesis whenever the p-value is smaller than 0.05. The empirical type I error is defined as the proportion of false rejections under each settings, as reported in Table S1.

### Evaluation of within-data structure preservation and batch effect removal

For the three negative control tasks Neg1-Neg3, and the five positive control tasks Pos1-Pos5, we compare the overall correlation between the pairwise distances of cells before and after alignment. Specifically, suppose 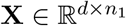 and 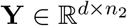 are the original normalized datasets, and 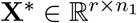 and 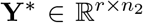 are the aligned datasets obtained by one of the integration methods, where *r* could be different from *d*. We first obtain 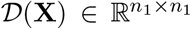, whose entries contain pairwise Euclidean distances between the columns of **X**. Similarly, we obtain *𝒟*(**Y**), *𝒟*(**X**^*∗*^), and *𝒟*(**Y**^*∗*^). Let [*𝒟*(**X**)]_*i*_ *b*e the *i*-th row of *𝒟*(**X**). We calculate Kendall’s tau correlations and Spearman’s rho correlations between [*𝒟*(**X**)]_*i*._ and [*𝒟*(**X**^*∗*^)]_*i*._, and between [*𝒟*(**Y**)]_*i*._ and [*𝒟*(**Y**^*∗*^)]_*i*._, for all *i*’s. The overall correlation associated with each integration method is then defined as the average correlation across all rows and across both datasets, as reported in the y-axis of Figures 2b, 3a and S5.

To evaluate the performance in batch effect removal using three different metrics. Specifically, once the integrated data are obtained, we calculate the Davies-Bouldin (D-B) index, and the inverse Calinski-Harabasz (C-H) index with respect to the batch labels. Specifically, for a given integrated dataset consisting of *K* batches *C*_1_, *C*_2_, …, *C*_*K*_, the D-B index is define as

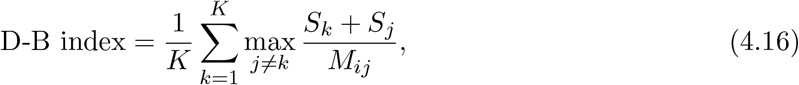

where

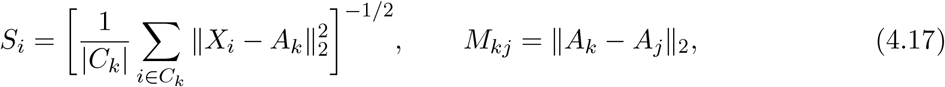

with *A*_*k*_ being the centroid of batch *k* of size |*C*_*k*_|, *X*_*i*_ is the *i*-th observed data point. The inverse C-H index is defined as

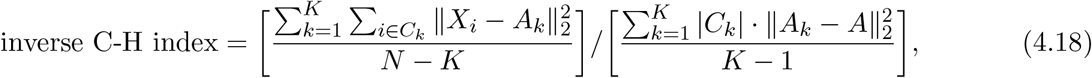

where *A* is the global centroid. As a result, we expect a method that achieves better alignment quality to have higher D-B index, and higher inverse C-H index. Compared with other metrics such as LISI and ARI, the above D-B and inverse C-H indices are directly calculated from the pairwise Euclidean distances, and do not rely on pre-specified nearest neighbor graphs (LISI), or the predicted cluster labels (ARI), which may be sensitive to specific methods.

### Simulation studies: DE gene detection

To further evaluate the performance of SMAI in identifying important marker genes, we generate datasets **X** ∈ ℝ^*d×n*^ and **Y** ∈ ℝ^*d×n*^, with *n* = 2100 and *p* = 1000, based on a Gaussian mixture model with variance *Σ*^*2*^ = 1 containing 12 clusters. Specifically, we first generate dataset **X**, so that each of its clusters has a mean vector uniquely supported on 20 features with the same value *ρ*. In other words, for each of the 12 clusters (cell types), there are 20 marker genes with nonzero mean expression levels within that cluster. Moreover, we consider unbalanced class proportions for **X**, by setting the cluster proportion as (0.18, 0.16, 0.15, 0.13, 0.11, 0.10, 0.03, 0.03, 0.03, 0.03, 0.03, 0.02). To generate **Y**, we first generate a dataset **Y**_0_ ∈ ℝ^*d×n*^ on the above Gaussian mixture model, but with a different cluster proportion (0.05, 0.05, 0.06, 0.07, 0.07, 0.08, 0.09, 0.09, 0.10, 0.11, 0.11, 0.12). Then we obtain **Y** by applying a similarity transform of **Y**_0_. In particular, we set 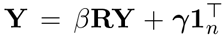, where *β* = 2, ***γ*** ∈ ℝ^*d*^ has random entries uniformly sampled from the interval [0, *ρ/*7], and **R** ∈ ℝ^*d×d*^ is a rotation matrix that slightly rotate the marker genes associated with Cluster 1.

To evaluate the performance of various alignment methods for assisting detection of marker genes, we apply each of them to obtain an integrated datasets **D** ∈ ℝ^*d×*2*n*^, concatenating the two datasets after alignment. Then, for each cluster, we perform one-sided two-sample t-tests with respect to each gene, to determine its associated marker genes. The sets of marker genes are obtained by selecting those genes whose p-values adjusted by the Benjamini-Hochberg procedure is below 0.01. Finally, for each cluster, we evaluate the agreement between the marker genes identified by each method, and the true marker genes. Specifically, we consider the Jaccard similarity index

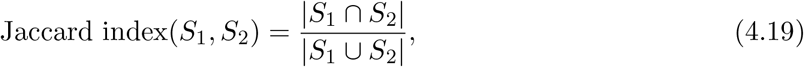

where *S*_1_ and *S*_2_ are two non-empty sets. In addition, we also assess the false positive rate

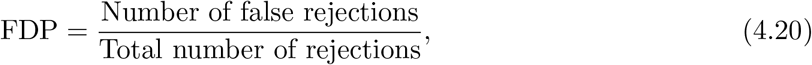

and the power

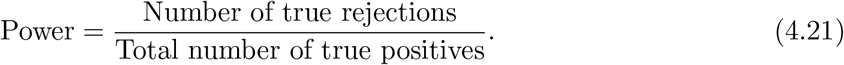

The simulation results corresponding to two different settings of {*ρ*, ***γ*, R**} are summarized in Figure S9.

### Differential expression analysis

For each of the positive control tasks Pos1-Pos5, we first obtain an integrated dataset containing all the samples, by using one of the alignment methods. Then for each cell type *k*, we identify the set of marker genes *S*_*k*_ based on two-sample t-tests, whose p-values are corrected by the Benjamini-Hochberg procedure. We use a threshold of 0.01 on the adjusted p-values to select the differentially expressed genes. In the meantime, for each cell type *k*, we also obtain the benchmark set 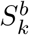 of marker genes, which contains all the differentially expressed genes identified based on the individual datasets before alignment. Finally, we compute the Jaccard similarity index between the set *S*_*k*_ of marker genes based on the integrated dataset, and the benchmark set 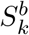 based on the original datasets. The results are reported in Figure 3c.

### Prediction of spatial genes

For the three spatial positive control tasks PosS1-PosS3, we withhold each gene from the spatial transcriptomic data, and predict its values based on the following procedure. In particular, we denote 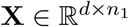 as the spatial transcriptomic data, and 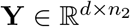 as the paired RNA-seq data. We also denote 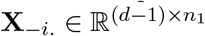 as the submatrix of **X** after removing the *i*-th row, and denote **X**_*i*._ ∈ ℝ^*d*^ as the *i*-th row of **X**. For each 1 *≤ i ≤ d*, we

1. Apply one of the alignment methods (LIGER, Scanorama, Seurat, fastMNN, or SMAI) to 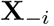 and 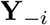, and obtain the aligned datasets 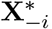 and 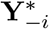 ;
2. Fit a *k*-nearest neighbor regression (with *k* = 5 in all our analysis) between the predictor matrix 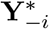 and the outcome vector 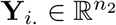;
3. Predict the outcome vector 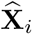. associated with the predictor matrix 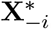 based on the above regression model.

To evaluate the accuracy of the predictions, we calculate the Kendall’s *τ* correlation between **X**_*i*._ and 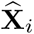.

### Computing time

We evaluate the computational time cost by SMAI-align with two existing alignment methods, Seurat and LIGER. Specifically, we consider a sequence of integration tasks involving datasets of various sample sizes. Seurat and LIGER are applied under the default tuning parameters, whereas SMAI-align is applied with *t* = 60 and *T* = 5. As a result, we find SMAI-align has a similar running time as Seurat, and is in general much faster than LIGER (Figure S13a). In particular, to align two datatsets with sample size *n*_1_ = *n*_2_ = 7000 and *d* = 2000, it took at most 7 mins on a standard PC (MacBook Pro with 2.2 GHz 6-Core Intel Core i7) for SMAI to obtain the similarity transformation and the aligned datasets. On the other hand, we find that even after including the SMAI-test procedure, the complete SMAI algorithm still has a reasonable computational time, that is, about twice of the time for running SMAI-align alone. For example, it takes less than 13 mins in total to align datasets each containing 7000 samples and 2000 features.

### Synthetic data

Synthetic datasets are generated to evaluate the performance of SMAI. Specifically, new pairs of datasets associated with the positive control Task Pos2 and the negative control Task Neg1 are generated by selecting subsets of cells from the datasets associated with the positive control Task Pos1. For Task Pos2, we generate the first dataset by removing the *beta* and *endothelial* cells from the Smart-seq2 dataset in Task Pos1, and generate the second dataset by removing the *gamma* cells from the CEL-Seq2 dataset in Task Pos1. As a result, the datasets in Task Pos2 have partial overlapping cell types. Similarly, for Task Neg1, the first dataset is such that it only contains the *acinar, activated sttellate, alpha* and *endothelial* cells of the Smart-seq2 dataset in Task Pos1, whereas the second dataset only contains the *beta, delta, ductal* and *gamma* cells of the CEL-Seq2 dataset in Task Pos1. The datasets in Task Neg1 thus do not share any common cell types.

### Data preprocessing

The raw counts data from each dataset listed in Table 1 were filtered, normalized, and scaled by following the standard procedure (R functions `CreateSeuratObject`, `NormalizeData` and `ScaleData` under default settings) as incorporated in the R package `Seurat`. For datasets with large numbers of genes, we also applied the R function `FindVariableFeatures` in `Seurat` to identify the top 2000 most variable genes for subsequent analysis. For the human pancreatic data associated with Task Pos1, we removed the cell types containing less than 20 cells. For the lung data associated with Task Neg3, a subset of 4000 cells were randomly sampled from each of the original datasets for our analysis.

### Implementation details

For all the integration tasks analyzed in this study, we use the R implementations of the relevant alignment algorithms. For LIGER and Harmony, we set the dimension required for the dimension reduction step to be 50. The other tuning parameters of LIGER and Harmony, as well as those of Seurat, Scanorama, and fastMNN, are set as their default values. We have implemented our SMAI-test and SMAI-align algorithms into an unified R/Python function, where under the default setting, we have *T* = 5, *t* = 60, and *k* = 20. To obtain the initial alignment for partial alignability test, we use Seurat, fastMNN, or SMAI-align to align **X**_2_ and **Y**_2_. We find the results to be not sensitive to the choice of these methods. In our analysis, we used the above default values for *T, t* and *k*, and used (4.14) for determining the rank parameter, with c = 0.001 for SMAI-align and c = 0.5 for SMAI-test. The R package “SMAI,” along with its Python version, and the R codes for reproducing the analyses and results presented in this study, can be retrieved and downloaded from our online GitHub repository https://github.com/rongstat/SMAI.

## Data Availabiliy

The human pancreatic data can be accessed in the R package `SeuratData` [https://github.com/satijalab/seurat-data] under the dataset name `panc8`. The PBMC data can be accessed in the R package `SeuratData` [https://github.com/satijalab/seurat-data] under the dataset name `pbmcsca`. The mouse brain chromatin accessibility data were downloaded from Figshare [https://figshare.com/ndownloader/files/25721789], containing a dataset from Fang et al. [29] (single-nucleus ATAC-seq protocol), and a 10X Genomics dataset (sample retrieved from [https://support.10xgenomics.com/single-cell-atac/datasets/1.2.0/atac_v1_adult_brain_fresh_5k]). The human lung data were downloaded as Anndata objects (samples 1, A3, B3 and B4) on Figshare [https://figshare.com/ndownloader/files/24539942]. The human liver and MLN data were downloaded from https://www.tissueimmunecellatlas.org. The mouse gastrulation seqFISH data were downloaded from https://content.cruk.cam.ac.uk/jmlab/SpatialMouseAtlas2020/, and the RNA-seq (10X Chromium) data can be accessed as ‘Sample 21’ in `MouseGastrulationData` within t he R package `MouseGastrulationData`. For the mouse VISP data, the ISS spatial transcriptomic data can be downloaded from https://github.com/spacetx-spacejam/data, the ExSeq spatial transcriptomic data can be downloaded from https://github.com/spacetx-spacejam/data, and the Smart-seq data can be downloaded from https://portal.brain-map.org/atlases-and-data/rnaseq/mouse-v1-and-alm-smart-seq.

## Code Availabiliy

The R and Python packages of SMAI, and the R codes for reproducing our simulations and data analyses, are available at our GitHub repository https://github.com/rongstat/SMAI.

## Supplementary Materials to

### 1 Supplementary Results

#### Simulations demonstrate empirical consistency and statistical validity of SMAI

To verify the statistical validity and consistency of SMAI, we generate two families of noisy datasets, each containing a distinct low-dimensional structure as its underlying true signal. Under each setting, a pair of (alignable) datasets are generated whose underlying low-dimensional structures are identical up to a similarity transformation. More specifically, for a given sample size *n*, we generate *d*-dimensional noisy data matrix **Y ∈**ℝ^*d×n*^ from the signal-plus-noise model **Y** = **Y**^*∗*^ + **Z**, where the columns of **Y**^*∗*^ are the underlying noiseless samples (signals), and **Z** is the random noise matrix. The signal matrix **Y**^*∗*^ are generated such that its columns 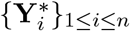 are sampled from some low-dimensional structure isometrically embedded in the *d*-dimensional Euclidean space. Each low-dimensional structure lies in some *r*-dimensional linear subspace, and is subject to an arbitrary rotation in ℝ^*d*^, so that 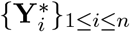 sare generally *d*-dimensional vectors with dense (nonzero) coordinates. The entries of **Z** are generated independently from a standard Gaussian distribution *N* (0, 1). Similarly, we generate another noisy data matrix **X** ∈ ℝ^*d×n*^ from 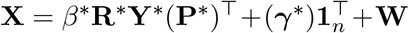, for some fixed scaling factor *β*^*∗*^ ∈ ℝ, mean shift vector ***γ***^*∗*^ ∈ ℝ^*d*^, permutation matrix **P**^*∗*^ ∈ ℝ^*n×n*^, rotation matrix **R**^*∗*^ ∈ ℝ^*d×d*^, and random noise matrix **W** ∈ ℝ^*d×n*^, containing independent standard Gaussian entries. Observe that both datasets **X** and **Y** are based on the same low-dimensional noiseless signal matrix **Y**^*∗*^. In this way, we have simulated noisy datasets **X** and **Y**, each having an intrinsically *r*-dimensional structure, that are identical up to a similarity transformation encoded by the parameters (*β*^*∗*^, ***γ***^*∗*^, **R**^*∗*^). Note that there is an one-to-one correspondence between (*β*^*∗*^, ***γ***^*∗*^, **R**^*∗*^) and the parameters in the optimization (2.1). We consider a smiley face with *r* = 2 (Figure S2a) and a Swiss roll manifold with *r* = 3 as the true signal structures (Figure S2b), where the signals 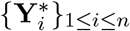 are sampled uniformly from each object.

For SMAI-align, we assess the quantitative errors (Methods) of the SMAI-inferred similarity transformation parameters 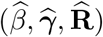, obtained from Algorithm 2, as compared with the underlying true transformation parameters (*β*^*∗*^, ***γ***^*∗*^, **R**^*∗*^), respectively, for various *n*. For SMAI-test, we evaluate the empirical type I errors against the above null model at the *α* = 0.05 level. Each setting is repeated for 300 times to obtain the empirical evaluation. We observe that for both signal structures the estimation errors decrease as *n* increases (Figure S2c), suggesting the consistency of the SMAI estimators 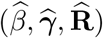 in the large-sample limit. In addition, the empirical type I errors (Table S1) are close to the nominal level *α* = 0.05 across all the settings, demonstrating the statistical validity of the test.

**Table S1:** Empirical type I errors of SMAI-test at the nominal level *α* = 0.05 under various signal structures and sample size *n*, with *d* = 1000.

| Low-Dimensional Structure | $n = 1200$ | $n = 1600$ | $n = 2000$ | $n = 2400$ |
| --- | --- | --- | --- | --- |
| Smiley Face | 0.062 | 0.063 | 0.040 | 0.043 |
| Swiss Roll | 0.043 | 0.046 | 0.040 | 0.043 |

### 2 Proof of Theorem 4.1

We begin by providing more details of the model assumptions. Throughout, we assume *r* is fixed and does not grow with *n* or *d*, indicating the low-dimensionality of the underlying structure. Without loss of generality, we also assume *β* = 1, as the discussion below can be easily adapted to allow for a global rescaling of all the eigenvalues of **Σ**_1_, or **Σ**_2_. In this case, under the null hypothesis, we are essentially interested in determining if 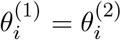 for all 1 *≤ i ≤ r*. Specifically, for the bulk eigenvalues 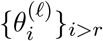, we assume that

**(A1)** The empirical distribution 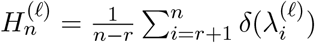 converges to a limiting distribution *H*^(*ℓ*)^ as *n → ∞*, where *δ*(*x*) denotes the Dirac mass at a point *x*.

For the assumption on the spiked eigenvalues 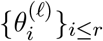, we define for any *α ≠* 0 lying outside the support of *H*^(*ℓ*)^ the following function

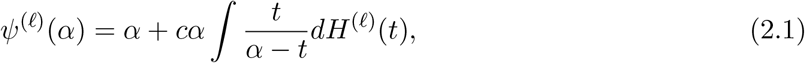

and its derivative

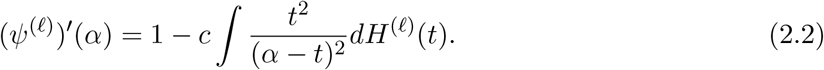

Similarly, we can define 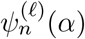 and 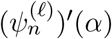, with *H*^(*ℓ*)^ in (2.1) and (2.2) replaced by 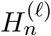, and *c* replaced by *n/d*. Then we further assume that

**(A2)** For each = 1, 2, the spiked eigenvalues 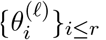 lie outside the support of *H*^(*ℓ*)^, and satisfy 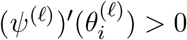 for all 1 *≤ i ≤ r*.

Finally, we also assume that the empirical bulk eigenvalues 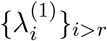 and 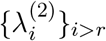 have similar empirical distributions. Specifically, we assume that

**(A3)** For each *i* = 1, 2, …, *r*, it holds that

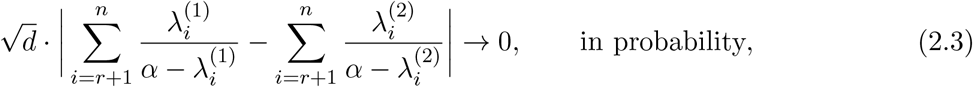

for all 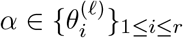

With these preparations, we proceed to prove our theorem. Firstly, we recall the following result obtained by Zhang et al. [2].

#### Lemma 2.1.

*for each* = 1, 2, *we define* 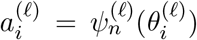. *Under the above high-dimensional generalized spiked covariance model and assumptions (A1) and (A2), it holds that*

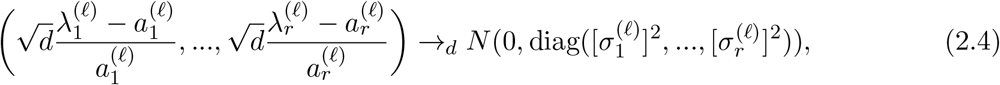

*where* 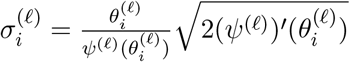

Now for any *i* = 1, 2, …, *r*, we have

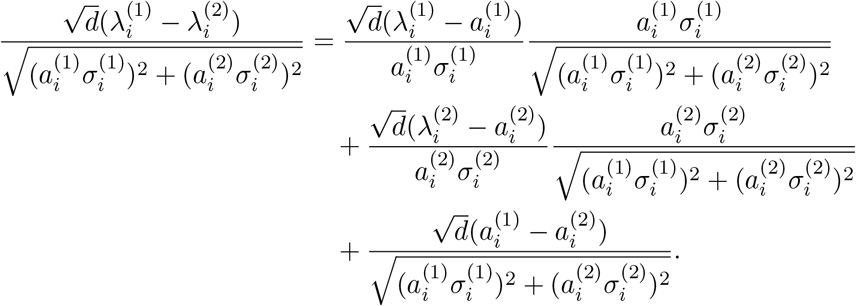

By Lemma 2.1, it follows that

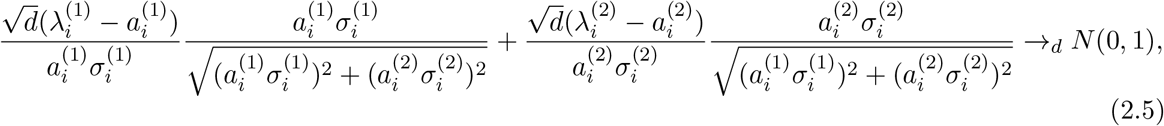

whereas by Assumptions (A1) and (A3), it follows that

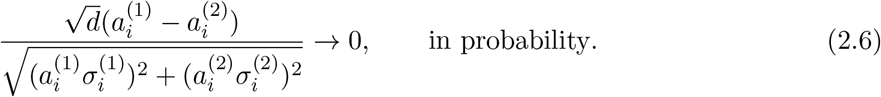

As a result, by Slutsky’s theorem, we have

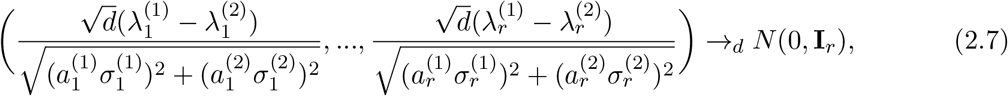

or

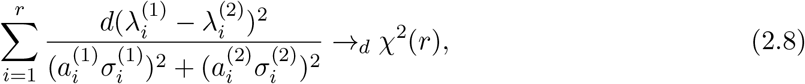

where *χ*^2^(*r*) is a *χ*^2^ random variable with degree of freedom *r*. Finally, in light of Algorithm 1, it suffices to show that

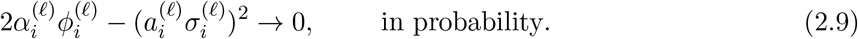

Note that by definition, we have

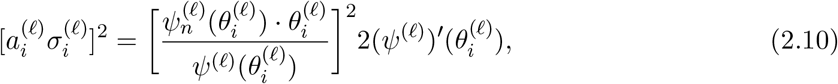

where

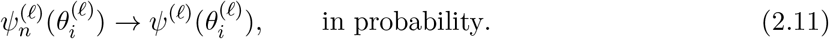

By Bai and Ding [1], we have

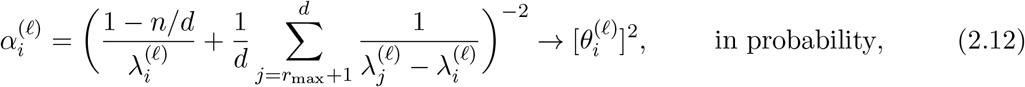

whereas by Section 2.4 of Zhang et al. [2], we have

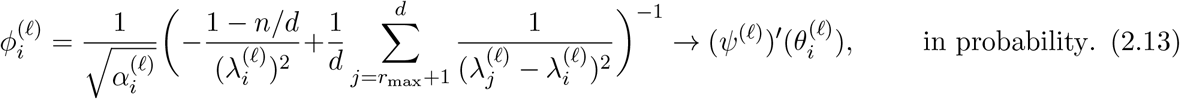

Combining Equations (2.10) to (2.13), we have shown Equation (2.9). This completes the proof of the theorem.

## Supplementary Figures

**Figure S1:**
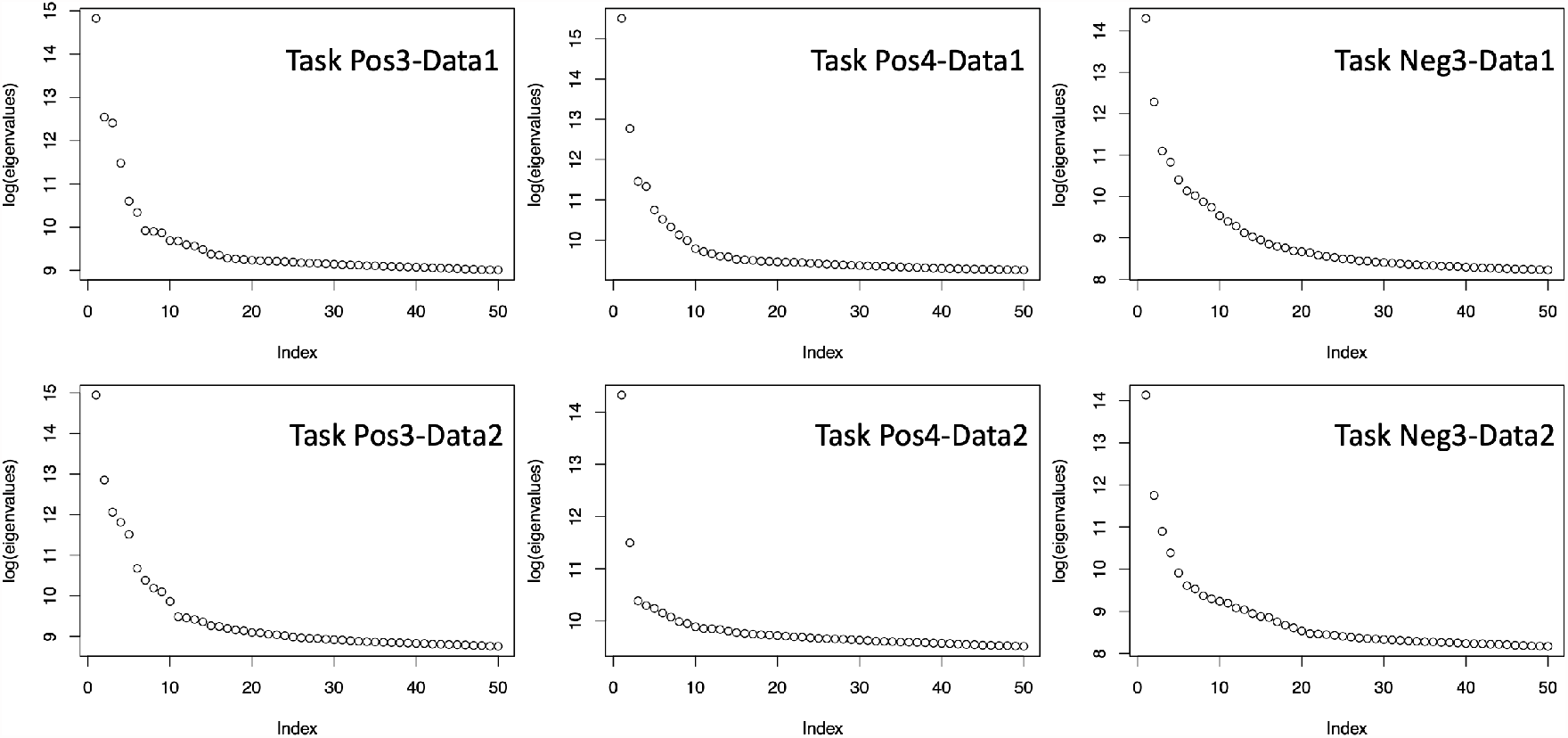
Empirical evidence on spiked eigenvalues. The top 50 eigenvalues associated to the normalized count data involved in some integration tasks are shown in order, at the log-scale. The presence of a few leading eigenvalues that are much larger than the rest of the eigenvalues indicates suitability of the spiked population models adopted by SMAI.

**Figure S2:**
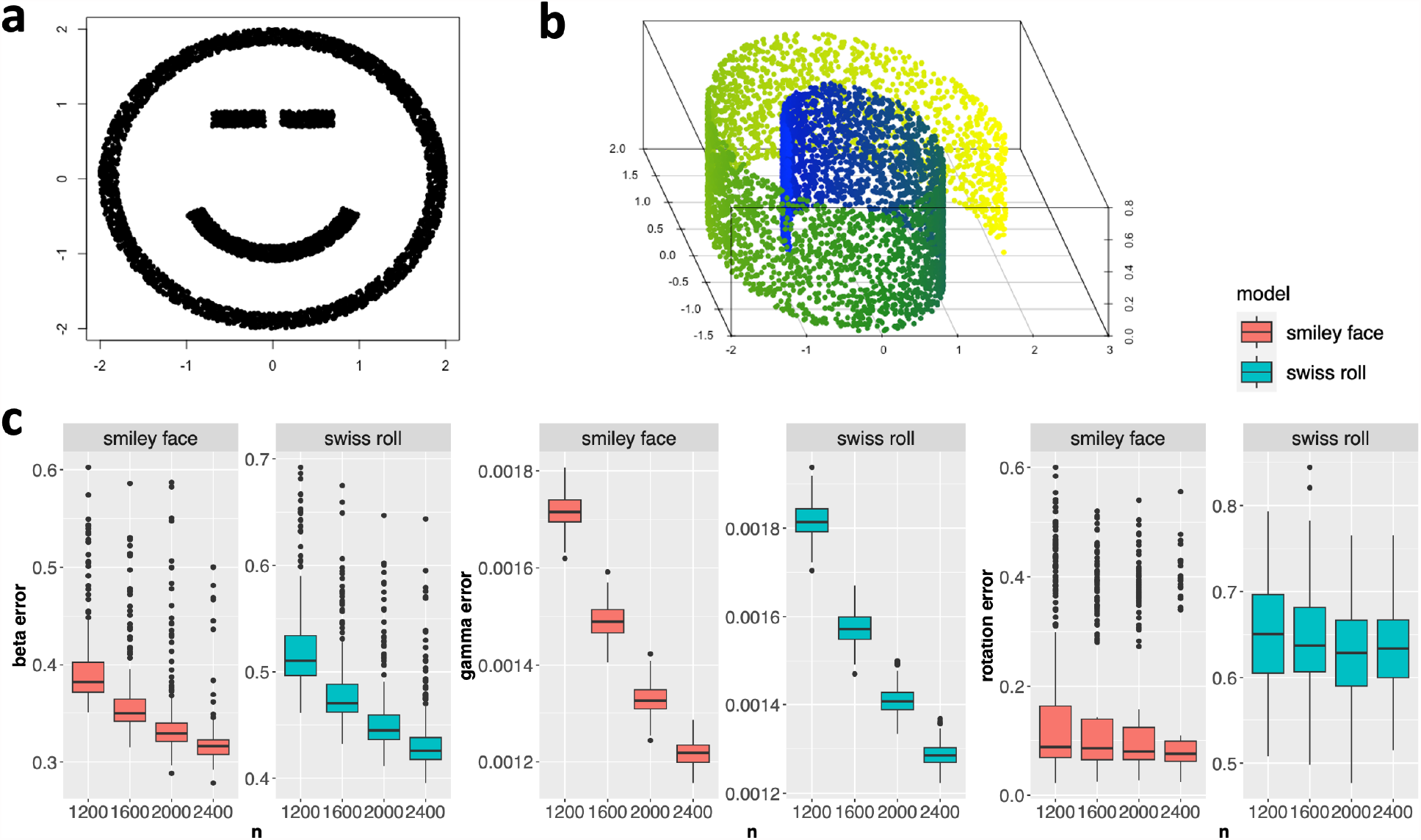
Results from simulation studies that demonstrate consistency of SMAI-align. (a) Illustration of *n* = 5000 samples from the two-dimensional smiley face structure. (b) Illustration of *n* = 5000 samples from the three-dimensional Swiss roll manifold. (c) Boxplots (center line, median; box limits, upper and lower quartiles; points, outliers) of estimation errors for *β*^*∗*^ (left), ***γ***^*∗*^ (middle), and **R**^*∗*^ (right) by SMAI-align algorithm, under the simulated data with two different underlying structures and various sample sizes *n*. Each boxplot is based on 300 rounds of simulations. The estimation errors decrease as *n* increases, suggesting the consistency of SMAI-align in recovering the true alignment function.

**Figure S3:**
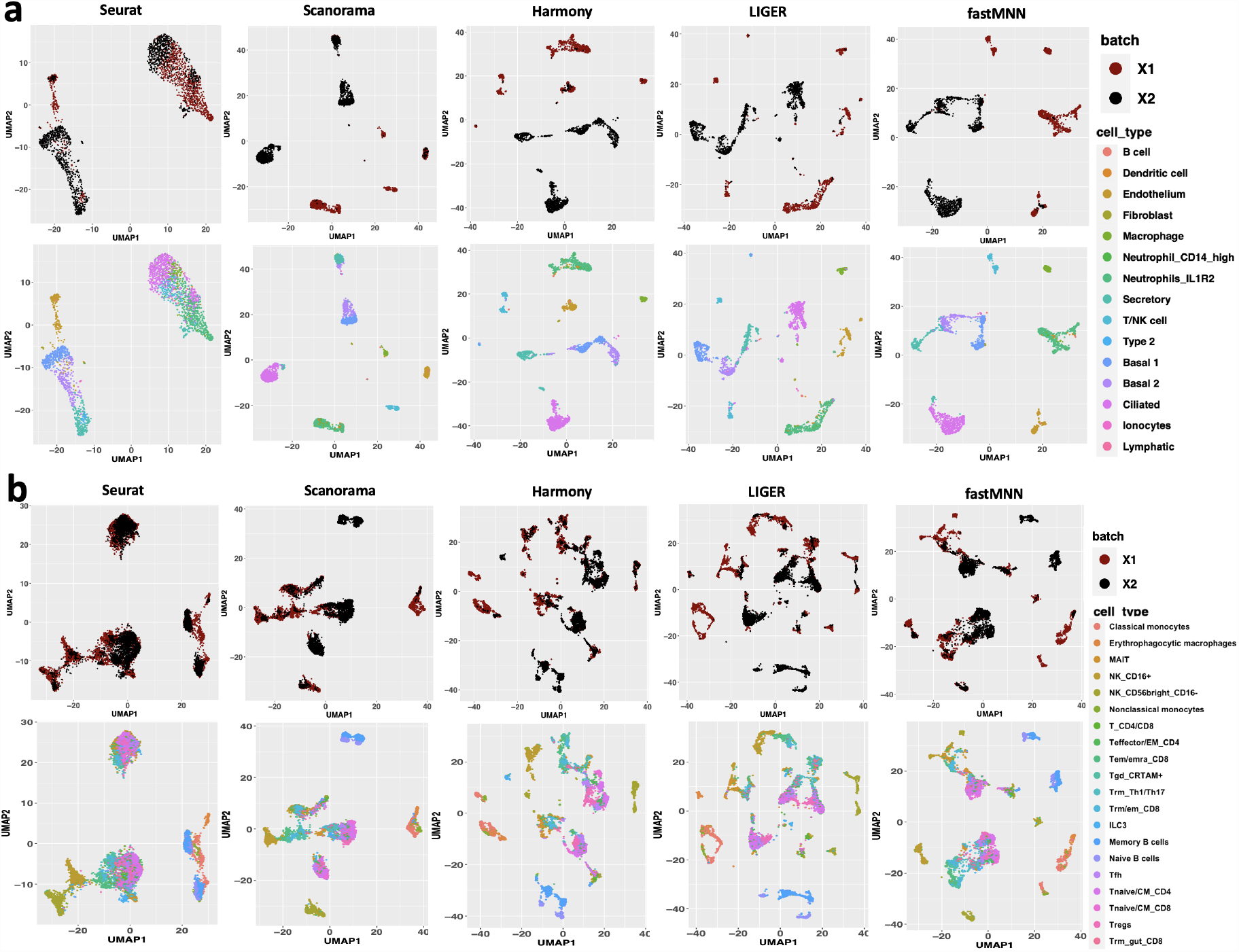
Evidence of distortion and misleading alignment from enforced data integration of negative control tasks. (a) and (b): UMAP visualizations of the integrated data under Task Neg2 (a) and Task Neg3 (b), as obtained by five popular methods. For each method, the top figure is colored to indicate the distinct datasets being aligned, whereas the bottom figure to indicate different cell types.

**Figure S4:**
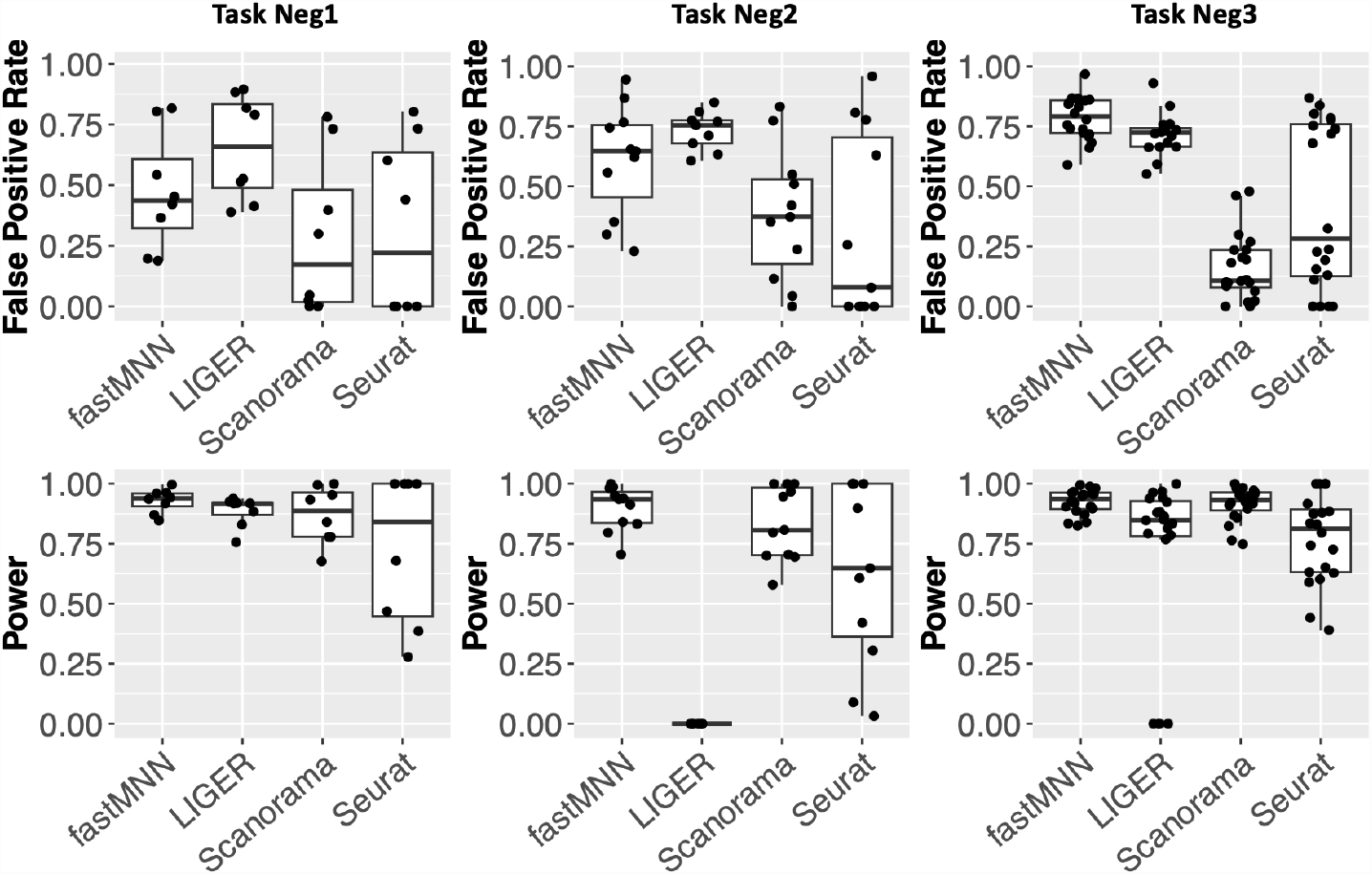
Effects on DE analysis caused by enforced data integration of negative control tasks. Boxplots of false positive rate and power of detecting DE genes based on the integrated data, as compared with the DE genes detected based on the original data. The distortion introduced by enforced alignment of two unalignable datasets may cause false discoveries and reduced power in detecting DE genes.

**Figure S5:**
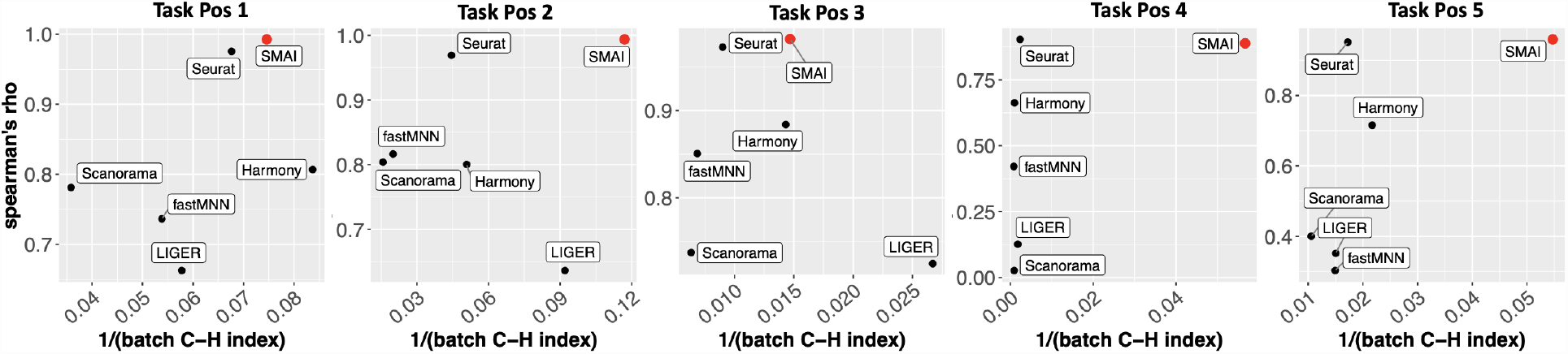
Evidences of improved structure-preserving integration by SMAI. Compared with five existing algorithms (black), SMAI-align (red) has uniformly better performance in preserving the within-data structures after integration while achieving overall the best performance in removing the unwanted variations. The structure-preserving quality is measured by Spearman’s rho correlations (y-axis), and the alignment quality is measured by inverse batch-associated C-H index (x-axis). For both metrics, a higher value suggests better performance.

**Figure S6:**
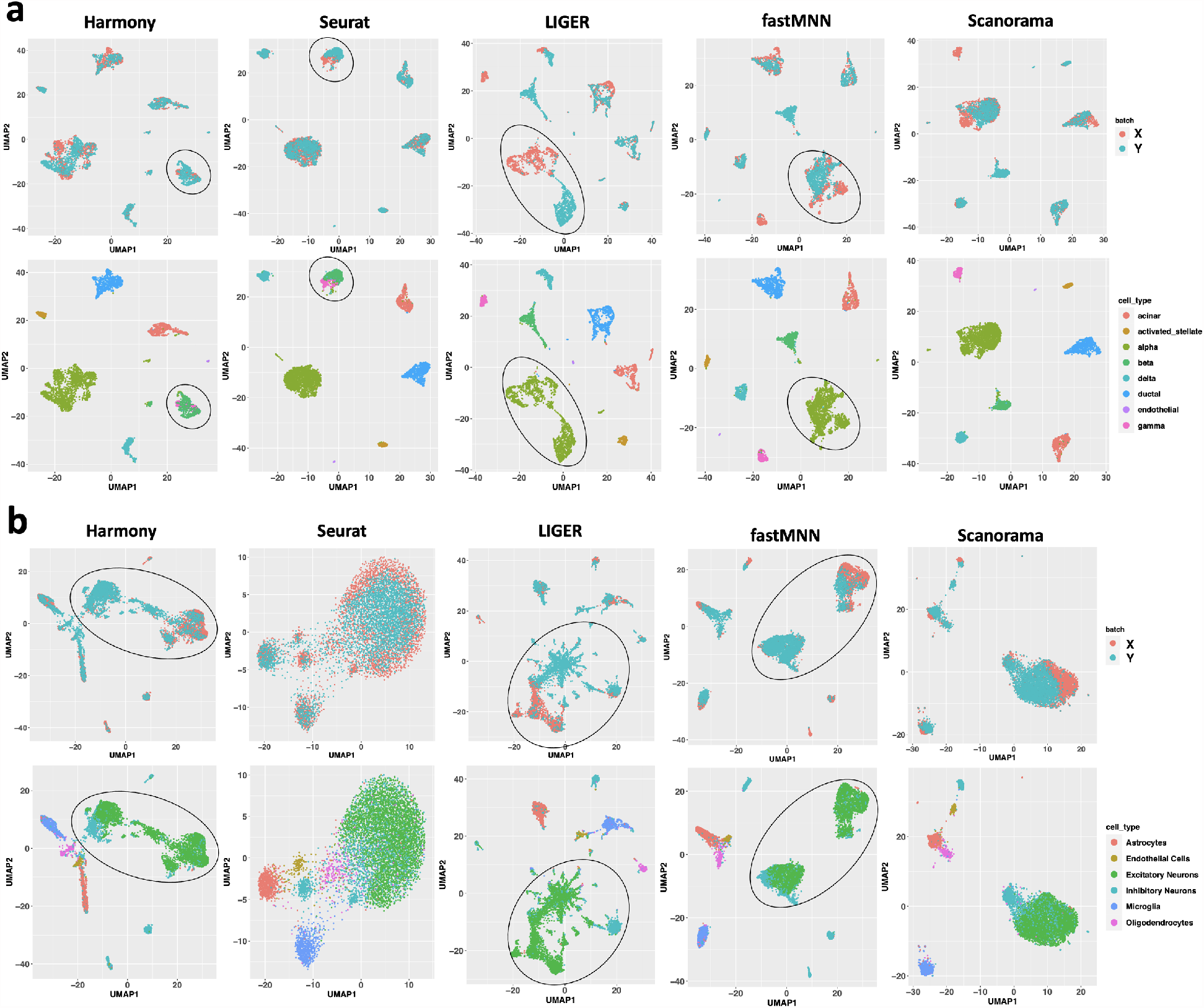
Distortion and misalignment by existing methods over positive control integration tasks. (a) In Task Pos2, there is false integration of gamma cells and beta cells by Harmony and Seurat, and significant distortion, that is, stretching and creation of multiple artificial subclusters, of the alpha cell cluster by LIGER and fastMNN. (b) In Task Pos4, there is strong distortion and artificial clustering of the excitatory neurons and the inhibitory neurons, in the data output from Harmony, LIGER and fastMNN.

**Figure S7:**
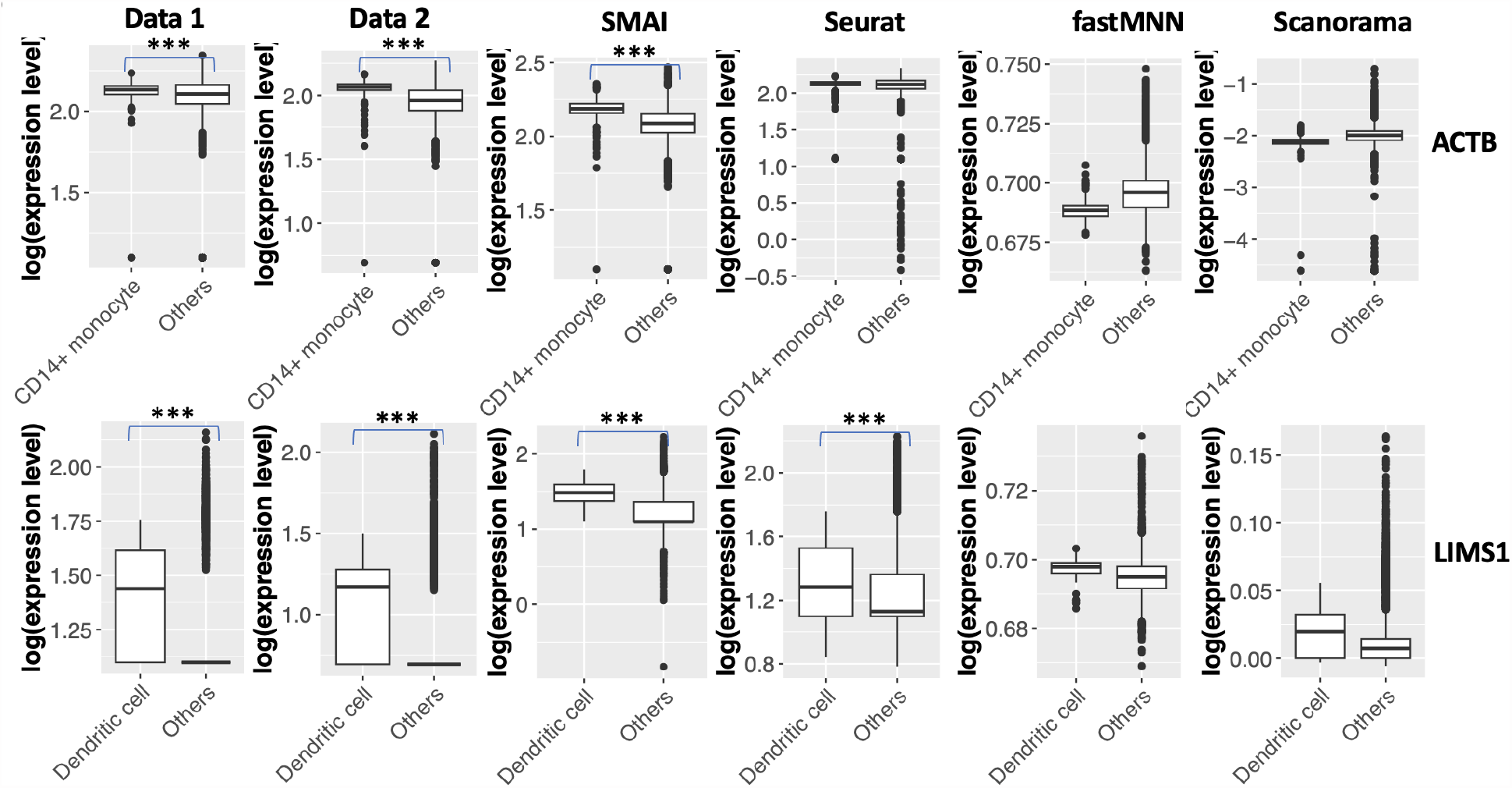
Examples about SMAI improving power of DE analysis. Top: Boxplots of expression levels of ACTB as grouped by cell types in the two datasets (Data 1 with 436 CD14+ monocytes and 2926 other cells, and Data 2 with 354 CD14+ monocytes and 2868 other cells) about human PBMCs (Task Pos3), and in the integrated datasets (790 CD14+ monocytes and 5794 others) as produced by SMAI-align, Seurat, fastMNN, and Scanorama. Bottom: Boxplots of log-expression levels of LIMS1 as grouped by cell types in the two datasets (Data 1 with 76 dendritic cells and 3286 other cells, and Data 2 with 38 dendritic cells and 3184 other cells) about human PBMCs (Task Pos3), and in the integrated datasets (114 dendritic cells and 6470 other cells) as produced by SMAI-align, Seurat, fastMNN, and Scanorama. In both examples, the DE pattern of the gene is not preserved by fastMNN, Scanorama, and/or Seurat, after integration. The stars above the boxplots indicate statistical significance of the p-values. Specifically, * means adjusted p-value *<* 0.05; ** means adjusted p-value *<* 0.01; *** means adjusted p-value *<* 0.001.

**Figure S8:**
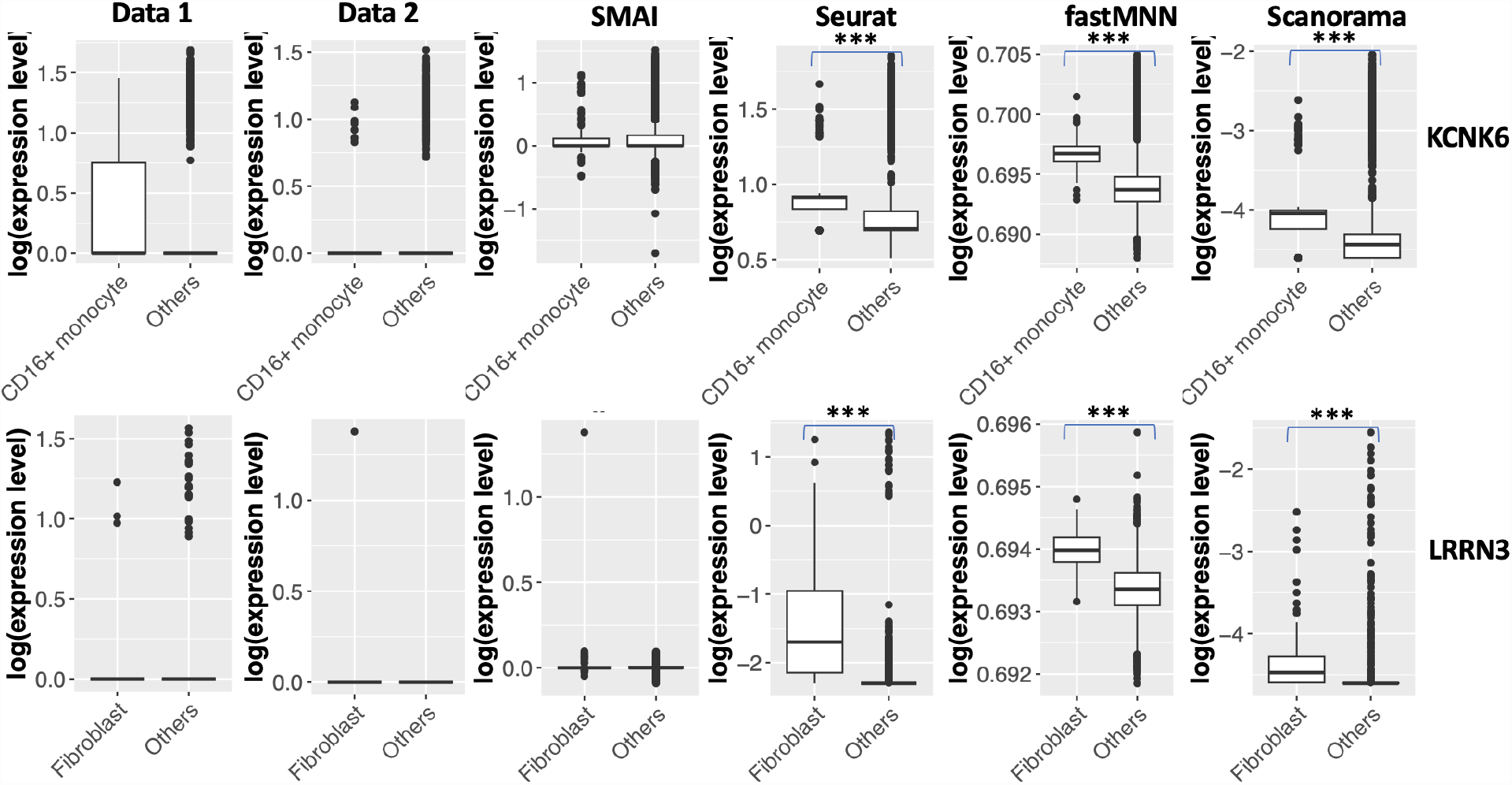
Examples about SMAI improving reliability of DE analysis. Top: Boxplots of expression levels of KCNK6 as grouped by cell types in the two datasets (Data 1 with 50 CD16+ monocytes and 3312 other cells, and Data 2 with 98 CD16+ monocytes and 3124 other cells) about human PBMCs (Task Pos3), and in the integrated datasets (148 CD16+ monocytes and 6436 others) as produced by SMAI-align, Seurat, fastMNN, and Scanorama. Bottom: Boxplots of log-expression levels of LRRN3 as grouped by cell types in the two datasets (Data 1 with 23 fibroblast cells and 2330 other cells, and Data 2 with 124 fibroblast cells and 1787 other cells) about human lung tissues (Task Pos5), and in the integrated datasets (147 fibroblast cells and 4117 other cells) as produced by SMAI-align, Seurat, fastMNN, and Scanorama. Artificial DE patterns are created by Seurat, fastMNN, and Scanorama after integration. The stars above the boxplots indicate statistical significance of the p-values. Specifically, * means adjusted p-value *<* 0.05; ** means adjusted p-value *<* 0.001; *** means adjusted p-value *<* 0.0001.

**Figure S9:**
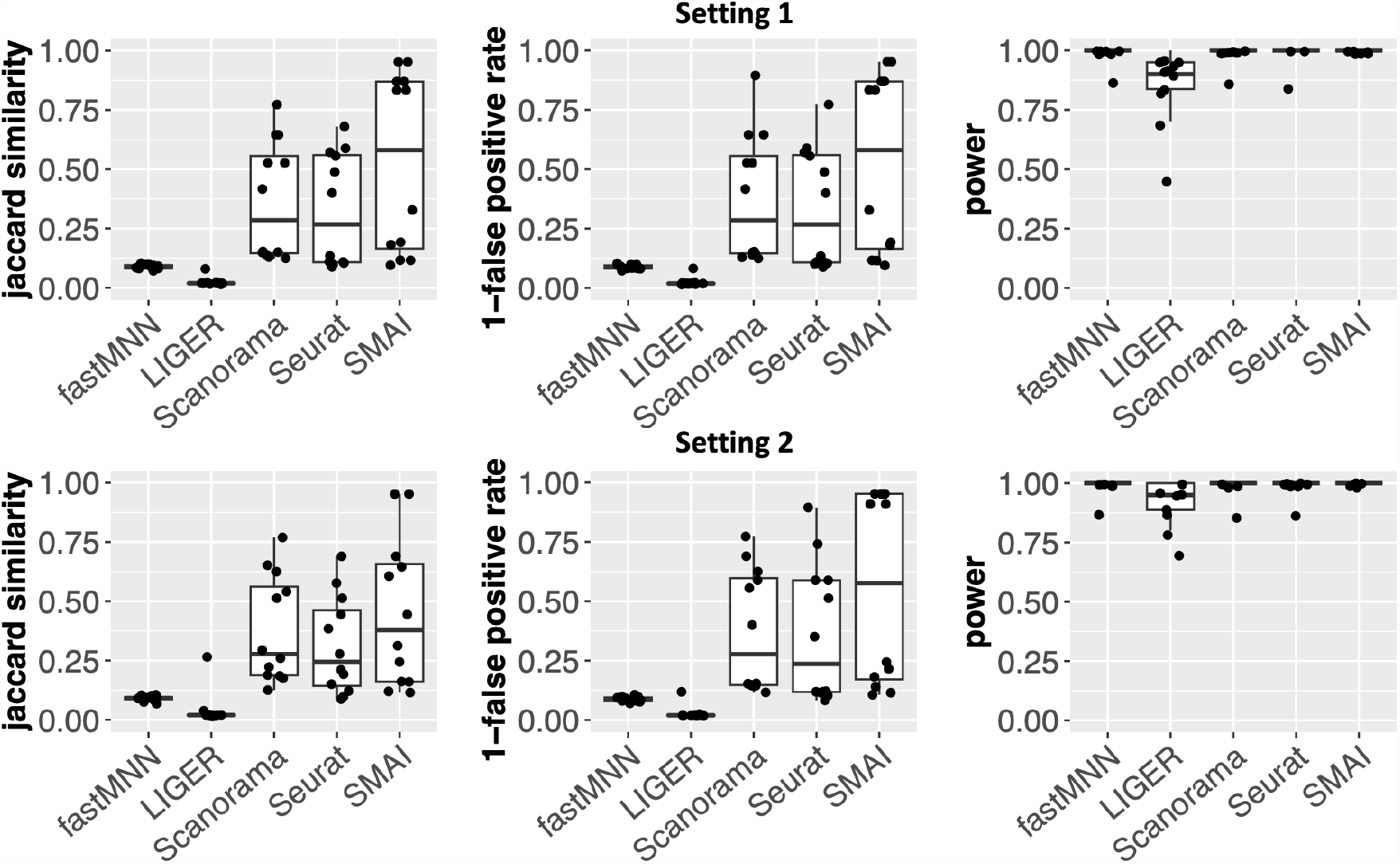
Simulation study reveals the advantage of SMAI in improving DE analysis. We carry out simulation studies by generating pairs of datasets, each containing about 2000 cells of 12 different cell types (clusters) and expression levels of 1000 genes, but with some batch effects and slight differences in cell type proportions. For the two different settings of signal-to-noise ratios, we show boxplots of the Jaccard similarity, the (1*−* false discovery rate), and the power, of the subsequent DE analysis with respect to the true marker genes, based on one of the integrated datasets as obtained by the five methods. Our simulation shows better performance of SMAI in precisely characterizing the true marker genes as measured by all the three metrics. In particular, our results suggest the tendency of including more false positives by the existing methods than SMAI.

**Figure S10:**
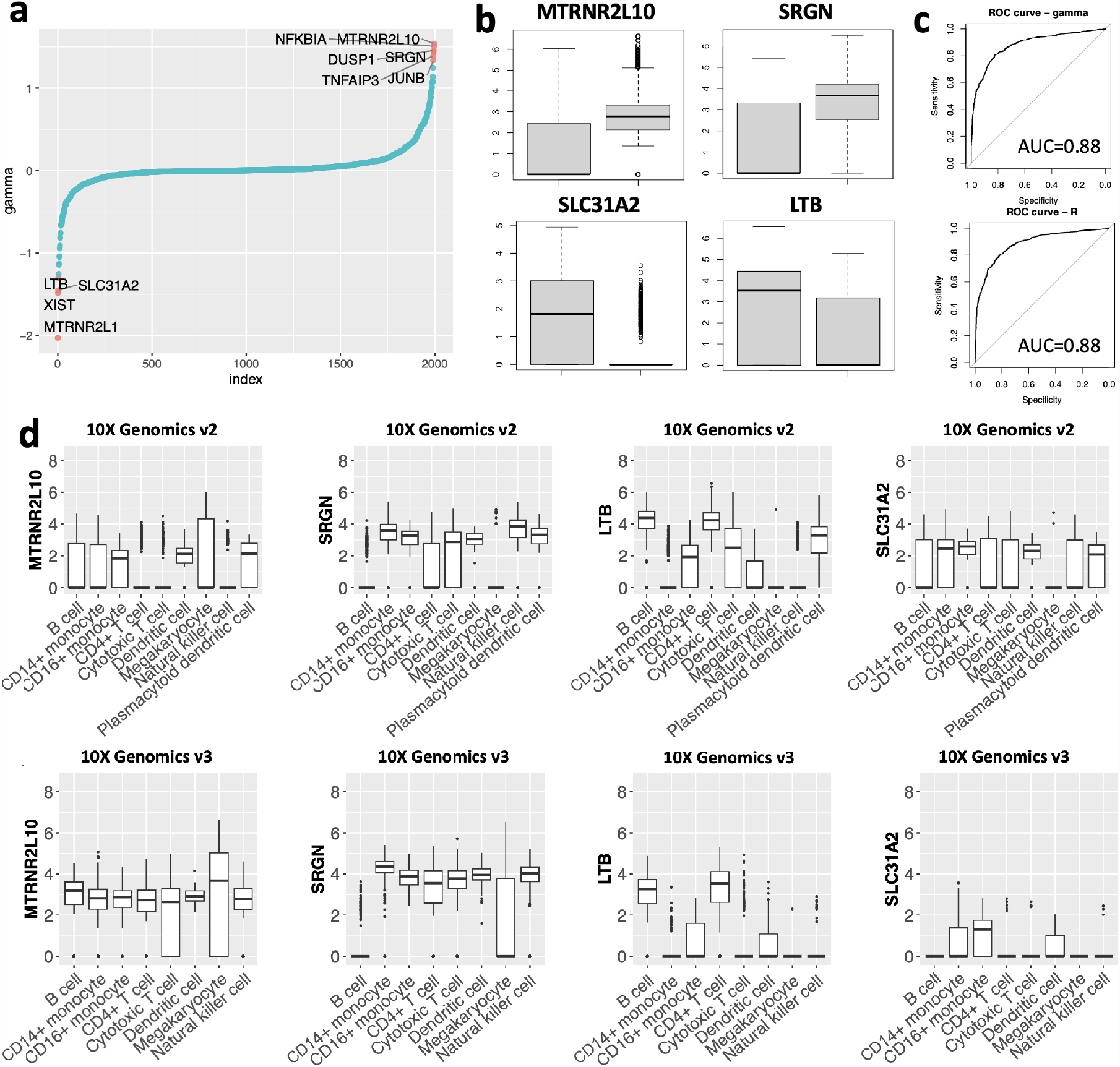
Additional evidences on SMAI’s interpretability and insights on the batch effects. (a) Visualization of the estimated mean-shift vector 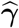 sfrom integrating the human PBMCs data (Task Pos3), whose components are ordered from the smallest to the largest. (b) Boxplots of a few genes with largest absolute values in 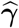 shown in (a). Left: 10X Genomics v2 dataset with *n* = 3362. Right: 10X Genomics v3 dataset with *n* = 3222. (c) ROC curves between the genes differentially expressed in one of the cell types in the PMBCs data (Task Pos3), and the batch-effect-related genes as captured by 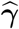 (Top), or by 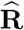 (Bottom). The AUCs suggest most of the genes contributing to the batch effects are simultaneously DE genes associated with certain cell types. (d) Boxplots of the genes from (b) and group by different cell types. The main discrepancy between the measured expression profiles across datasets is likely caused by the differences in the total counts of these transcripts under the respective sequencing technologies.

**Figure S11:**
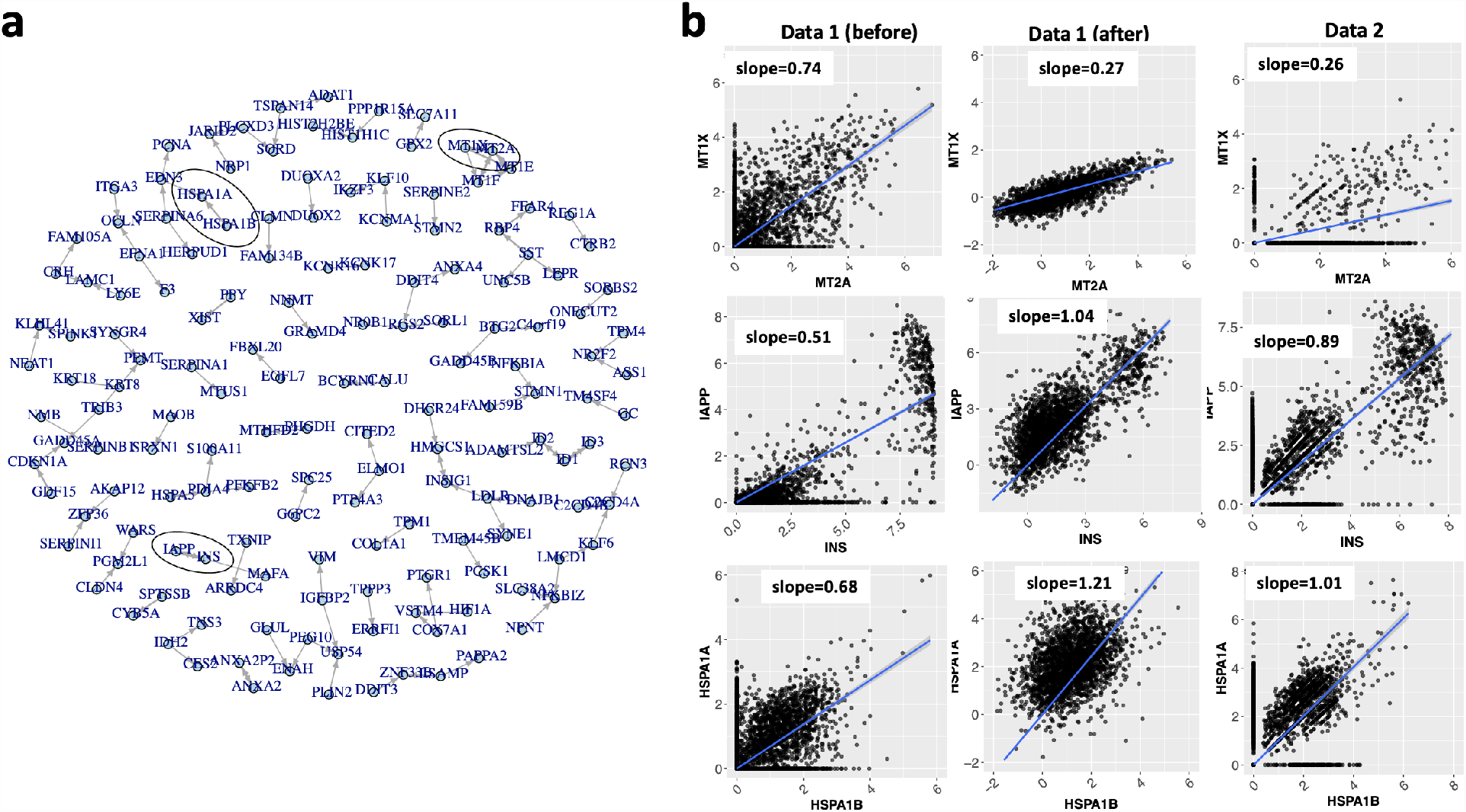
Insights on batch effects revealed by rotation 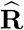 in Task Pos1. (a) A directed weighted graph representation of the rotation matrix 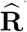 after removing the edges with weight *<* 0.1. The genes linked by thicker edges are more significantly altered during alignment. (b) Scatter plots and the slope of fitted lines between the expression profiles of gene pairs highlighted in (a). Left: measurements in dataset 1 (Smart-Sea2) before alignment to dataset 2 (CEL-Seq2). Middle: measurements in dataset 1 after alignment to dataset 2 by SMAI. Right: measurements in dataset 2. SMAI-align leads to more similar co-occurrence patterns between the genes, as quantified by the slopes, in the two datasets after data alignment.

**Figure S12:**
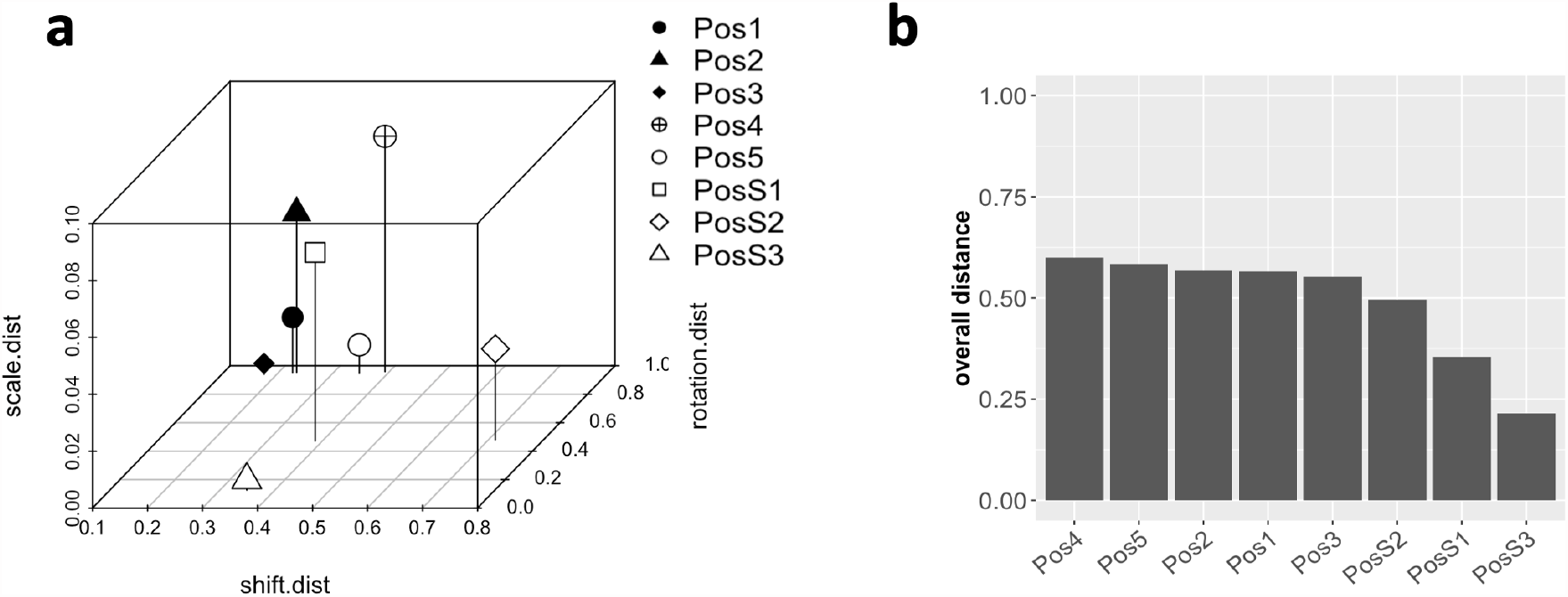
SMAI enables geometric quantification of batch effects. (a) 3-D scatter plot of the batch effects across all the positive control tasks, as characterized by the obtained SMAI-align parameters 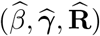, converted into three distance metrics between 0 and 1, with larger values indicating relatively greater amount of rescaling (z axis, or scale.dist), translation (x axis, or shift.dist), or rotation (y axis, or rotation.dist) needed to achieve alignment. (b) Barplot showing the overall distance between the aligned datasets, as characterized by an average of the three geometric distances shown in (a). These metrics enable quantifying the magnitude and the geometric composition of the underlying batch effects between the two datasets.

**Figure S13:**
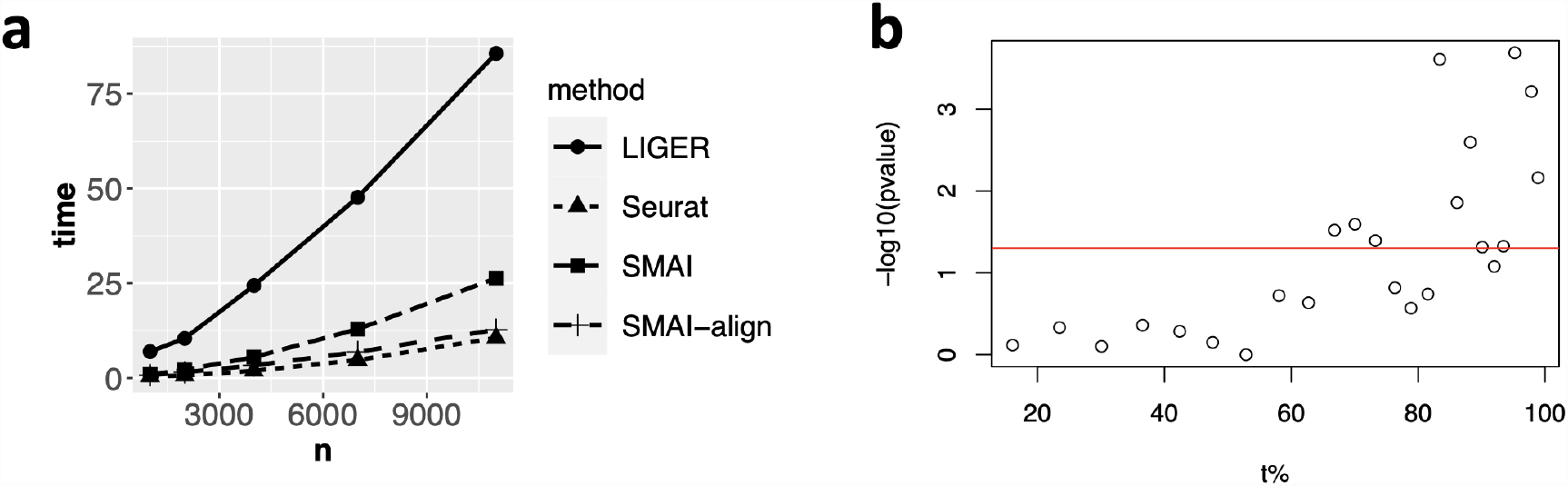
Computing time and effect of thresholding parameter. (a) By evaluating computing time over a series of datasets with improving sample sizes, we find SMAI-align has a similar running time as Seurat, and is in general much faster than LIGER. The complete SMAI algorithm including SMAI-test and SMAI-align still has a reasonable computational time, that is, about twice of the time for running SMAI-align alone. (b) Scatter point between the threshold *t* and the log-transformed p-values from SMAI-test concerning task Pos5. The red line corresponds to p-value = 0.05. When *t* is below certain value (e.g., 60%), the p-values are not significant as the null hypothesis about partial alignailibity is likely true; when *t* gets larger, the null hypothesis are mostly rejected. The transition point is likely the true proportion of alignable samples.

